# Actionable biological programs to enhance EGFR-targeted therapy response unveiled by single-cell lineage tracing in clinically relevant lung cancer models

**DOI:** 10.1101/2025.05.19.654081

**Authors:** Beatrice Gini, Whitney Tamaki, Johnny Yu, Dora Barbosa, Wei Wu, Yashar Pourmoghadam, Paul Allegakoen, Donghwa Kim, Sohit Miglani, Sarah Elmes, Victor Olivas, Hani Goodarzi, Trever Bivona

## Abstract

Developing high-resolution approaches to capture both tumor architectural clonality and transcriptional state(s) in individual cells within heterogeneous tumor cell populations could shed light on the evolution of pre-existing and newly emergent tumor subclones and their phenotypes, elucidating their trajectories in response to selective pressures such as drug treatment. Reports to date have focused primarily on analyzing the drug-induced evolution of lung cancer cells in *in vitro* preclinical models with limited complexity and a relative lack of characterization of actionable biological programs to induce durable responses. Here, we challenged this paradigm and employed a lineage tracing single-cell RNAseq method to track the evolution of primary non-small cell lung cancer (NSCLC) patient-derived organoids (PDOs) and tumor xenografts in response to the standard-of-care EGFR inhibitor osimertinib, with a focus on understanding drug persistence and resistance. Our single-cell lineage tracing-RNAseq system revealed the presence of a discrete set of lineages with distinct transcriptional phenotypes over the course of the treatment. We identified two lineage populations that became predominant during drug treatment and resisted therapy in the PDOs and tumor xenografts. These lineages were present before treatment and harbored Hedgehog pathway and FOXD1 transcriptional programs, respectively. These specific transcriptomic lineages were otherwise undetectable by lower-resolution profiling. Functional studies confirmed the protective role that the baseline expression of the Hedgehog pathway and FOXD1 programs in the lineage tumor cell sub-populations exerts upon targeted therapy. The potential clinical relevance of these regulatory programs was validated by cross-analysis of single-cell transcriptomic data obtained from human NSCLC specimens. Overall, our approach identified pre-existing seeds of resistance before therapy and convergent, adaptive mechanisms supporting tumor residual disease and resistant states. This study highlights the utility of high-resolution tracing of tumor clonal heterogeneity with matched single-cell profiling to reveal occult cell states and molecular mechanisms of therapy resistance and develop counteracting strategies.

## Main Text

Human cancers have highly heterogeneous genomic and transcriptomic landscapes with profound therapeutic implications^1–6^. Understanding the phenotypic consequences of the underpinning genomic heterogeneity concerning intrinsic or acquired targeted therapy resistance can inform more durable, tumor-specific therapies.

Advanced-stage Non-Small Cell Lung Cancer (NSCLC) is one of the most heterogeneous types of cancer, carrying high degrees of genomic and transcriptomic heterogeneity that impact both tumor progression and response to therapy^1,3,5–10^ and is one of the deadliest diseases in the United States, with a 5-year relative survival rate of 8% (American Cancer Society). Despite clinical successes with the use of molecularly targeted (and immune-based) therapies, drug resistance inevitably occurs either in the form of intrinsic or acquired resistance and results in lethal tumor progression^1,7,11^.

Among the main genetic subtypes of NSCLC, the Epidermal Growth Factor Receptor mutant (EGFRmt) cohort represents approximately 15% and 50% of lung adenocarcinomas in Western and Asian populations, respectively^11^. First-line osimertinib treatment has demonstrated notable efficacy compared to previous EGFR TKIs in untreated EGFRmt advanced NSCLC patients, achieving a significantly longer median progression-free survival (18.9 months with osimertinib vs. 10.2 months with standard EGFR TKIs) and longer median duration of response (17.2 months with osimertinib vs. 8.5 months with standard EGFR TKIs; FLAURA, phase III trial)^12^. Despite the improved clinical outcomes, responses to osimertinib are generally partial and incomplete as a result of drug persistence, and complete resistance (acquired resistance) almost inevitably subsequently occurs and limits long-term patient survival and cures^1,2^. Drug persistence and resistance can be caused by both genetic and, often, non-genetic mechanisms acting alone or in concert to overcome the drug treatment^2,13–15^.

Lineage tracing systems have been applied to investigate the evolution of various cancer models^16–19^. However, lineage tracing focusing on mechanisms of drug persistence in NSCLC models has been, to our knowledge, limited to conventional cell culture^16,20^. The detection of a pre-existent clonal population harboring a transcriptional state that confers resistance to treatment in more patient-proximate complex models could allow for the construction of more clinically relevant predictive models of the acquisition of drug resistance based on the *a priori* observation of that state within cancer cell populations. Therefore, we advanced on previous work and studied patient-derived primary organoids, which retain higher clonal heterogeneity than isogenic cell lines, better recapitulating clinical response to therapy^21,22^, and xenograft models to analyze the tumor evolution from treatment naïve to drug-persistent and resistant states.

We applied an ex vivo, static lentiviral barcoding system, matched with high-throughput, single-cell RNA sequencing (scRNA-seq), to track the evolution of transcriptionally-defined NSCLC clonal populations (lineages) at osimertinib-persistent and resistant states^5,11,23^, cross-validating the preclinical findings with unique clinical cohorts^5^ that contain scRNA-seq datasets obtained from EGFRmt clinical cases before and after EGFR-targeted therapy. This work is one of the first attempts to track osimertinib-induced tumor evolutionary trajectories in primary, patient-derived organoids and *in vivo* models with clinical validation, enabling biomarker discovery and enhanced therapeutic strategies for potential further clinical evaluation in the future.

### Results

## Analysis of the lineage evolution in primary EGFRmt NSCLC PDOs uncovers clonal populations with enhanced plasticity and persistence under osimertinib treatment

To expand the utility and accuracy of lineage tracing in organotypic and xenograft cancer models to enable an understanding of lineage-based changes across time in response to targeted therapy, we performed a study of NSCLC cancer cell lineage and cancer cell state by pairing genomic and scRNA-seq to simultaneously track the changes in clonal population frequencies and adaptations over treatment time (Fig. 1).

**Figure 1.**
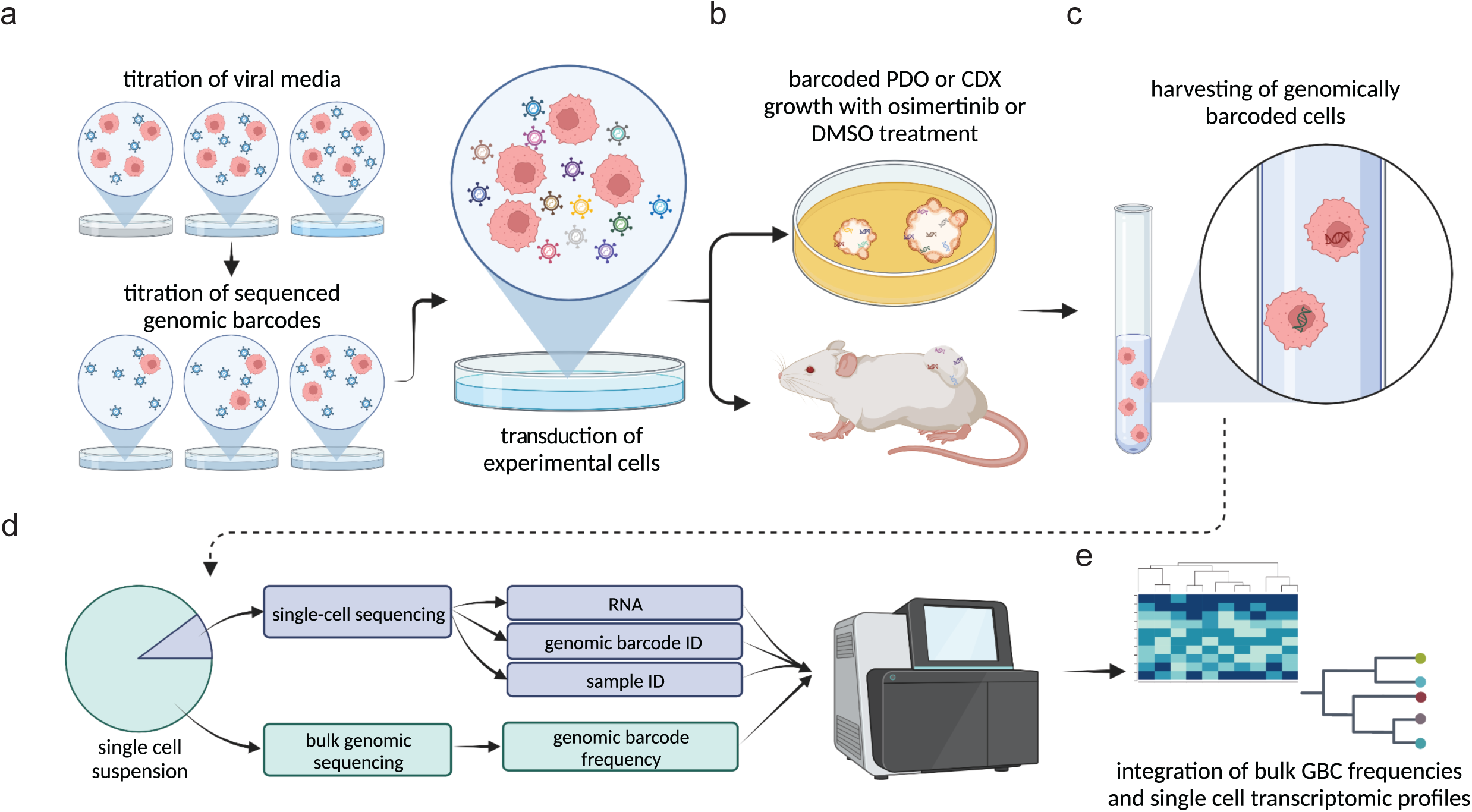
Static barcoding allows the generation of NSCLC models for scRNAseq analysis matched with lineage tracing: analysis workflow. (a) Barcoding of experimental cells through lentiviral transduction, (b) PDOs and CDXs establishment, (c) collection of single-cell suspensions for library preparations and sequencing (d), and (e) biocomputational analysis (see Methods).

After establishing an effectively barcoded founder population in our model systems (see Methods), we designed a study examining the effect of osimertinib over time. EGFR mutant NSCLC Patient-Derived Organoids (TH107 PDOs) were treated for up to 30 days with osimertinib or 14 days with DMSO control, and single cells and gDNA were collected at days 0, 7, 14, 21, and 30 (Fig. 2a and Supplementary Fig. 1a, b). Sensitivity to osimertinib was further characterized at day 19 (Fig. 2b) and day 37 (Supplementary Fig. 1c) to assess the sensitivity of EGFR mutant TH107 PDOs to osimertinib in this time frame. Despite the prolonged drug exposure (30 days), the organoids had reduced growth upon osimertinib treatment compared to treatment naïve parental cells at both time points and were accordingly classified as early persister TH107 PDOs at day 19 and late persister TH107 PDOs at day 30 (Supplementary Fig. 1d). Moreover, by the end of the osimertinib treatment, TH107 PDOs showed visible loss of organoid’s growth and 3D morphology, consistent with the effective drug treatment. Still, they were not eliminated, while the DMSO-treated organoids had rapid outgrowth by day 14, necessitating culture cessation (Fig. 2c).

**Figure 2.**
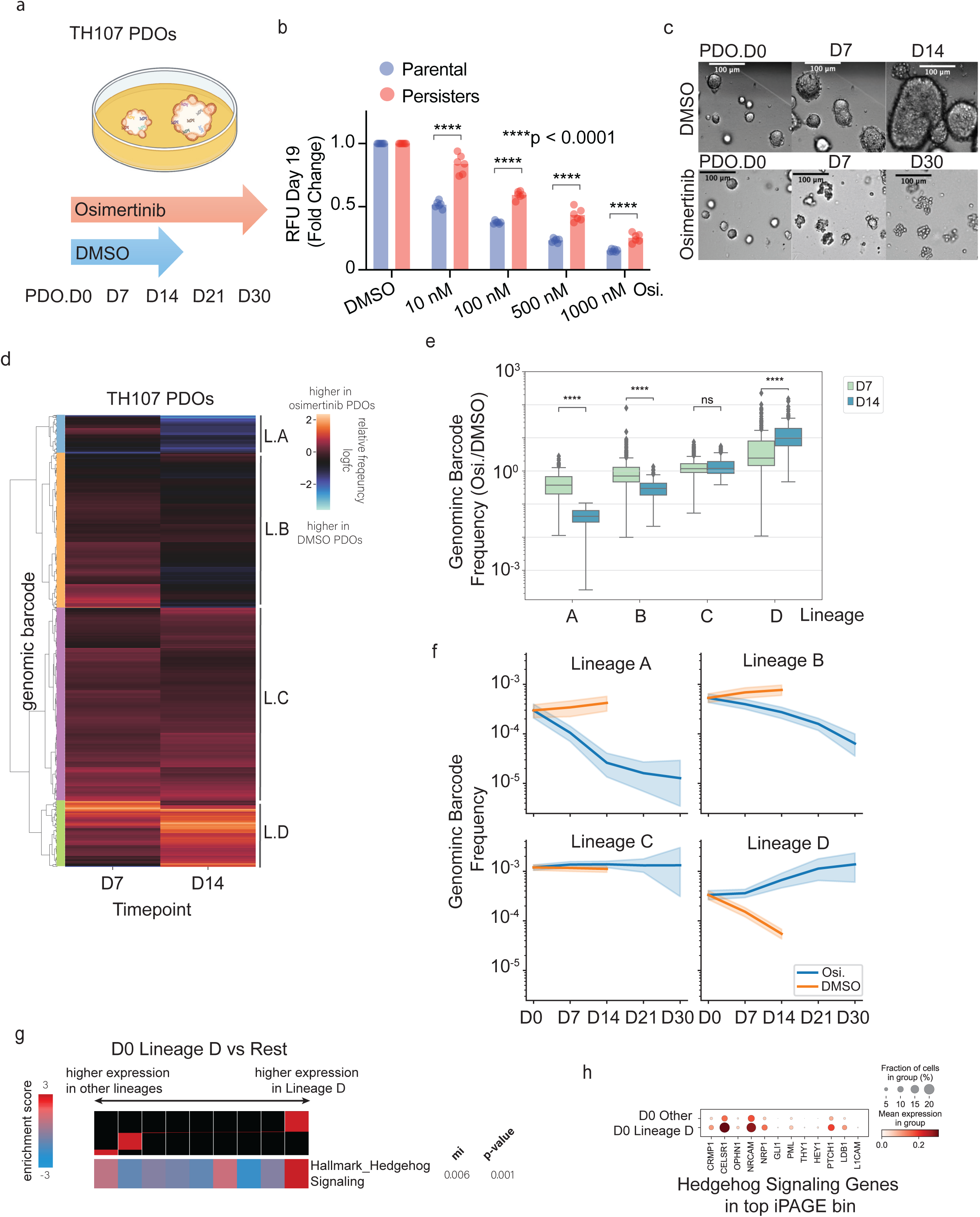
Analysis of the evolution and pathway gene expression of lineages detected in EGFRmt NSCLC TH107 PDOs reveal clonal populations persisting to EGFR inhibitor with upregulation of a subset of Hedgehog pathway genes in the treatment naïve state. (a) Time points of single-cell suspension collection from barcoded TH107 PDOs upon EGFR targeted therapy using osimertinib. (b, c) Assessment of osimertinib sensitivity by 3D-CTG (b) and high-magnification images (c) of TH107 PDOs at D0-D30 upon osimertinib or DMSO treatments (Parental: treatment-naive TH107 PDOs; Persisters: osimertinib persisters TH107 PDOs; p-value calculated using Two-way ANOVA with Sidak’s multiple comparison test). (d) Identification of four distinct lineage populations (L.A, L.B, L.C, L.D) differentially enriched at D7 and D14 in osimertinib-treated TH107 PDOs normalized to DMSO control. (e) Genomic barcode frequency scores for L.A-L.D populations highlight a gradual enrichment at D14 of the L.D population upon osimertinib treatment (p-values calculated using the Mann-Whitney-Wilcoxon test). (f) Lineage population abundances over the entire time course of the experiment show the distinctiveness behavior of the four lineage populations among osimertinb and DMSO-treated TH107 PDOs. (g) Pathway analysis of gene expression (iPAGE) comparing L.D population versus rest (L.A-L.C) at D0. All individual sequenced genes are sorted by log2FC and divided into 9 equally populated bins from lowest (negative) log2FC (left) to greatest log2FC (right). The red bar within the black header indicates the range of log2FC values in each bin. The heatmap is colored by the enrichment or depletion of the gene in each bin. The top/right-most iPAGE bin, containing genes with the greatest log2FC values, is significantly enriched for genes in the Hallmark Hedgehog Signaling gene set. The significance of the mutual information value (bits) is calculated using a non-parametric randomization test (see Methods). (h) Dotplot of Hallmark Hedgehog Signaling Gene Set genes present in the top iPAGE bin. See also Supplementary Figures 1, and 2.

To create a clean set of GBCs for integration with transcriptional data, we began with the 34,889 unique barcodes recovered from the T0 PDO sample. We then filtered for barcodes that occurred at least twice at T0 and were present at all other time points, leaving 2,384 barcodes to be overlaid onto the scRNA-seq data (Supplementary Fig. 2a). We performed unsupervised clustering of the normalized Genomic Barcode (GBC) frequencies across EGFR mutant TH107 PDO barcodes. We identified four major lineages with distinct population shifts after osimertinib treatment, compared to T0 (Fig. 2d and Supplementary Fig. 2a-c). Of these four lineages, only PDO Lineage D (PDO L.D) showed significant relative expansion in osimertinib-treated samples, normalized to DMSO-treated sample frequency, over time (Fig. 2e and Supplementary Fig. 2b). Furthermore, PDO L.D continued to expand in relative population frequency past D14, comprising 6% of the initial D0 population and increasing to 25% of cells at D30 (Fig. 2f and Supplementary Fig. 2b).

Having identified the lineage selected upon osimertinib treatment, we sought to understand transcriptional programs underlying survival under TKI treatment in this population. By overlaying lineage identity onto the matched scRNA-seq data, we investigated the transcriptional profile of the resistant lineage PDO L.D at D0. We extracted both pathway and gene-level change between L.D and all other lineages by leveraging iPAGE^24^ to perform gene set enrichment analysis (see Methods). iPAGE identified enrichment of Hallmark Hedgehog Signaling Pathway genes (Hh) in genes most upregulated in persister PDO L.D, compared to all other lineages, at D0 (Fig. 2g). Among the genes in this pathway, *CRMP1, CELSR1, OPHN1, NRCAM1, NRP1, GLI1, PML, THY1, HEY1, PTCH1, LBD1,* and *L1CAM* had the highest upregulation in PDO L.D compared to PDO L.A-L.C pretreatment (Fig. 2h). Among the top-upregulated genes *CELSR1, NRCAM1, NRP1, GLI1,* and *LBD1* are characterized tumor- and metastasis-promoting factors^25–29^, suggesting that these genes could be top candidate genes conferring an increased advantage to survival to osimertinib. Although previous studies reported that targeting the Hedgehog signaling synergizes with the EGFR signaling inhibition, ^30–32^ to our knowledge, this is the first association linking pre- treatment, higher expression of a subset of Hh pathway-related genes, to subsequent EGFR TKI drug persistence.

We compared the transcriptional profiles of PDO L.D at D30 to D0 using the iPAGE analysis to examine the changes in PDO L.D after treatment. iPAGE identified enrichment of genes within the Hallmark EMT gene set at D30 compared to D0, indicating that, over time, PDO L.D upregulated expression of EMT as a possible transcriptional adaptation to TKI treatment (Supplementary Fig. 2d-f). The EMT phenotype has been shown to promote EGFR-targeted therapy resistance in lung cancer^33,34^. Drug persister cells have been reported to undergo phenotypic changes to acquire stem-like properties, including expression of Hedgehog and EMT pathways, to enable a highly plastic persister state^35^.

Taken together, the data raise the possibility that high expression of Hh genes pretreatment may represent L.D’s increased propensity for phenotypic plasticity^36^, permitting the ability to undergo further transcriptional rewiring towards an EMT state during treatment over time that could confer increased survival to osimertinib compared to other lineages.

Identifying a set of cells expressing Hh pathway genes, pretreatment and expanding in relative population frequency over osimertinib persistence, was uniquely possible due to our paired static lineage tracing GBCs and scRNA-seq. We confirmed this by comparing the standard, single-cell RNA-sequencing pipeline to our single-cell RNA-sequencing matched with the lineage tracing method (Supplementary Fig. 3). Standard scRNA-seq analytic pipelines rely on the clustering of cells based on their gene expression profiles to group cells of similar transcriptional profiles for interpretable annotations^37^. Traditional clustering of the EGFR mutant TH107 PDO data, based only on gene expression profiles, identified six clusters (Supplementary Fig. 3a). Although Clusters 1 and 4 increased in relative population frequency over treatment, similar to L.D, none of the individual clusters expressed all the Hallmark Hedgehog Signaling genes identified by iPAGE (Fig. 2g, h, and Supplementary Fig. 3b, c). This supports the notion that our use of genomic barcodes for lineage tracing enabled the identification of PDO L.D as a potential sub-reservoir of cells that can rewire their transcriptional state to tolerate and resist EGFR-targeted therapy. Additionally, the static barcoding specifically enabled mapping a transcriptomic shift in PDO L.D D30 compared to D0. The signaling shift from high Hh to high EMT was undetectable when examining the bulk population (data not shown), likely due to the small proportion of PDO L.D. cells at D0.

Furthermore, we performed deep whole-exome sequencing on bulk EGFR mutant TH107 PDO samples to test whether the TKI treatment induced the selection or acquisition of *EGFR* gene alterations affecting therapy response. We did not detect additional *EGFR* mutations or copy number alterations in early and late persister-TH107 PDOs, compared to DMSO control (Supplementary Table 1), suggesting that acquired or pre-existing modifications of the EGFR target gene did not primarily confer the TKI-resistant phenotype observed in PDO L.D. This is consistent with the hypothesis that the higher expression of the Hh transcriptional program in PDO L.D pretreatment and enhanced lineage plasticity could confer osimertinib persistence.

## Pharmacological inhibition of the Hh pathway enhanced sensitivity to osimertinib in primary EGFRmt NSCLC PDOs

We sought a deeper functional characterization of the Hh transcriptional program identified in PDO L.D pretreatment, which could confer enhanced plasticity and osimertinib persistence. Although differential gene expression analysis did not highlight any significantly upregulated PDO L.D surface proteins at D0, limiting the possibility of purifying PDO L.D from bulk barcoded TH107 cells for additional analyses, we performed pharmacological inhibition of bulk EGFR mutant TH107 PDOs to test the hypothesis of whether inhibiting the Hh transcriptional program pharmacologically could increase sensitivity to osimertinib and prevent the transition of the population into the EMT state. We performed acute (5 days) and long-term (21 days) combinatorial treatment with osimertinib and the Hedgehog pathway inhibitor sonidegib^38^ in the EGFR-mutant PDOs (Fig. 3 and Supplementary Fig. 4). Sonidegib inhibits the Hh pathway by blocking the activity of the transmembrane protein Smoothened (SMO). When Hh signaling is absent, PTCH inhibits signaling by SMO. When Hh ligands bind to PTCH, the complex is internalized and degraded, enabling SMO accumulation and downstream activity^38–40^. Long-term combinatorial treatment resulted in significantly reduced organoid formation compared to single-agent therapy, using a low concentration range of osimertinib, suggesting the possibility of combining an Hh inhibitor to minimize potential toxicity induced by osimertinib in a clinical setting (Fig. 3a-c). Moreover, inhibition of the Hh pathway, which acts in synergy with the EGFR signaling^30^, in combination with osimertinib, more successfully inhibited proliferation and EGFR signaling compared to each monotherapy, with concomitant upregulation of apoptosis as measured by cleaved PARP levels (Fig. 3d), and reduction of the EMT biomarkers Claudin-1 and Slug (Fig. 3e, f).

**Figure 3.**
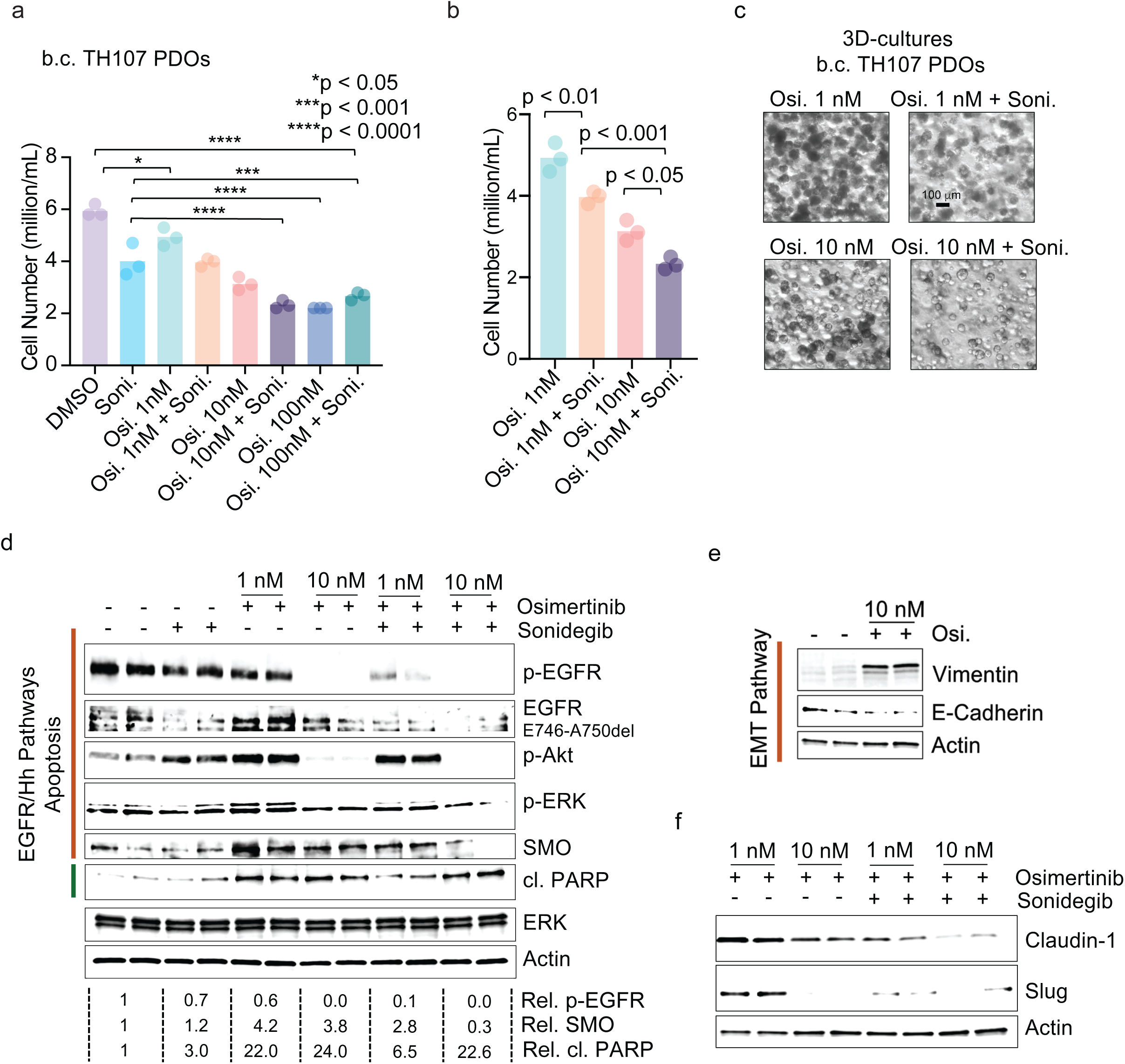
Combination therapy of Hedgehog and EGFR inhibitors rescues sensitivity to EGFR therapy in TH107 PDOs, pre-empting transition toward an EMT phenotype. (a-c) Pharmacological inhibition of Hedgehog pathway using sonidegib (Soni.) in combination with osimertinib in TH107 PDOs; PDO cultures were quantified (a, b) and imaged at the end of experiments (c). (d) Biochemical analysis of EGFR signaling, SMO expression, and apoptosis upon mono- and combinatorial treatment with sonidegib and osimertinib; average, fold change of p-EGFR, SMO, and cleaved PARP provided underneath each blot (band normalized to Actin and expressed as a fold change compared to DMSO treated controls). (e, f) Biochemical analysis of EMT biomarkers (Vimentin, E-Cadherin) upregulation upon osimertinib treatment (e) and EMT biomarker inhibition (Claudin-1, Slug) upon the combination with sonidegib (f) in TH107 PDOs. (n = 3 per experimental group; p-value calculated using one-way ANOVA and Tukey’s multiple comparison test; Soni.: Sonidegib, Osi.: Osimertinib). See also Supplementary Figure 4.

Our data indicate that combination therapy of Hh inhibitor with osimertinib can enhance response to EGFR inhibitor therapy in primary EGFRmt NSCLC organoids that contain occult cells that are potentially primed for transcriptional plasticity, EMT, and therapy persistence (Fig. 3d, e) and more potently suppresses the adaptation toward a therapy-persistent EMT phenotype (Fig. 3f). Thus, combination therapy could block the plastic potential of Hh-expressing cells and reduces their ability to undergo EMT or other adaptive transcriptional rewiring and increase sensitivity to osimertinib. The ability to identify and target a plasticity primed state provides a different approach to address therapy resistance from more conventional strategies.

## Analysis of the lineage evolution *in vivo* in EGFRmt NSCLC CDXs uncovers osimertinib-resistant clonal populations with an upregulated FOXD1 program

To better capture the *in vivo* context of the response to osimertinib in NSCLC and ensure the generalizability of our platform, we generated barcoded EGFRmt H1975 Cell-Derived Xenografts (CDXs), which were then treated with osimertinib (5mg/kg) for 32 days (Fig. 4a and Supplementary Fig. 5a, b) (see Methods). Single cells were collected at four time points along the course of the TKI treatment: the first time point of collection, at D0, allowed us to characterize the representation of uniquely barcoded clonal populations before TKI treatment; D12 represented a persistence state, whereas tumor growth resumed by D18, and thus D18 and D32 were selected as early and late resistance states, respectively (Fig. 4a).

**Figure 4.**
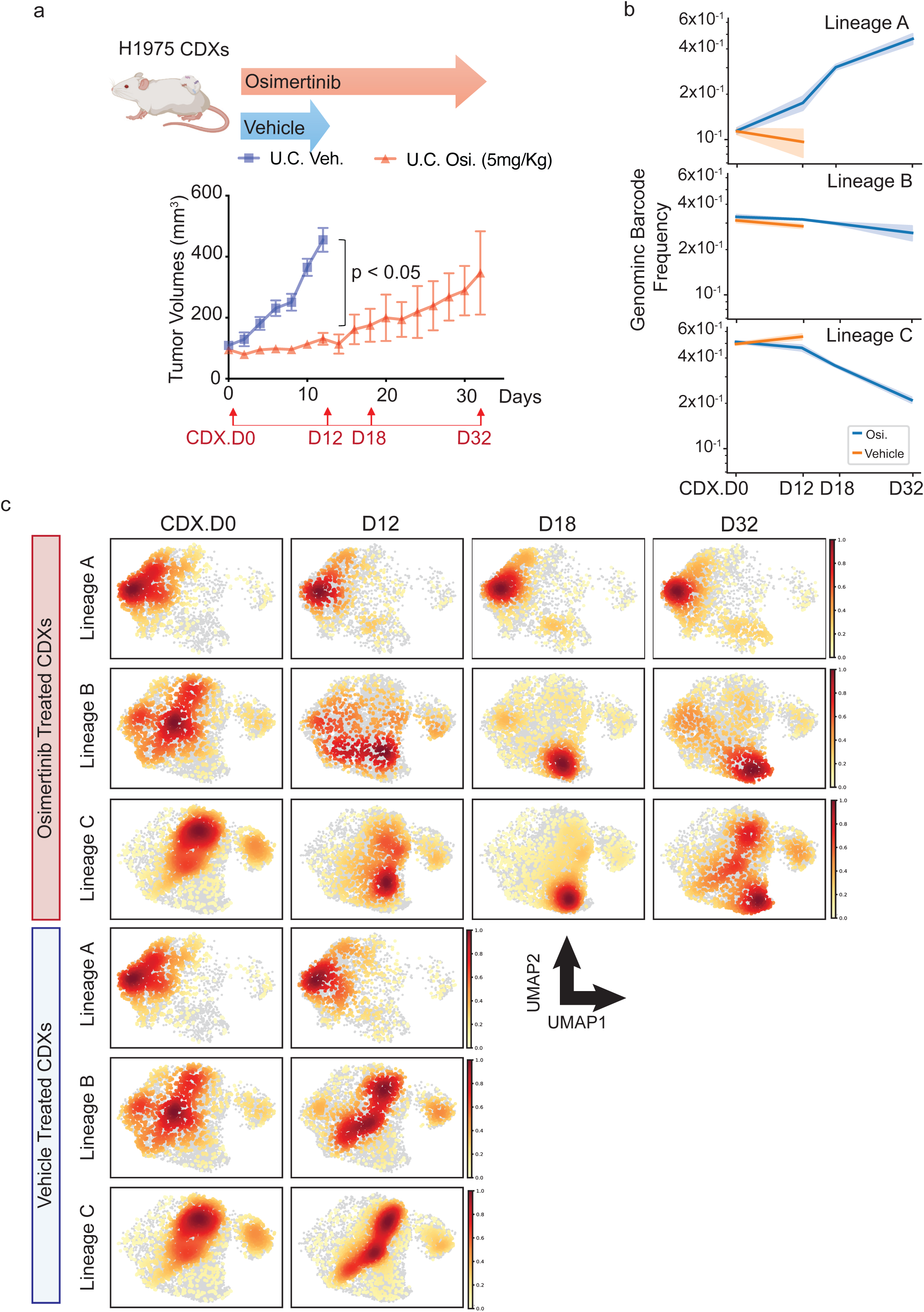
Analysis of the evolution of lineages detected in EGFRmt NSCLC CDXs uncovers three lineage populations with different sensitivity to osimertinib treatment and distinct transcriptional signatures. (a) Growth curve of vehicle and osimertinib treated barcoded H1975 CDXs and corresponding timeline of lineage tracing matched with scRNAseq analysis (p-value calculated using Student T-test). (b) Identification of three distinct lineage populations (A, B, C) differentially enriched at D0 and D12-32 in osimertinib-treated H1975 CDXs compared to vehicle control (b). (c) UMAP projections of lineage population A, B and C in osimertinib and vehicle treatment groups over the course of the entire in vivo experiment (D0-D32) showing distinct transcriptional phenotypes. See also Supplementary Figure 5.

Next, we applied our lineage tracing analysis pipeline to study the transcriptomic transitions of the tumor lineage populations along the pretreatment and drug persistent to resistant states in our CDX EGFRmt H1975 NSCLC models, complementing the PDO studies of osimertinib persistence.

GBC clustering for H1975 CDX cells identified three lineages with distinct transcriptional signatures and relative abundance over time (Fig. 4b and Supplementary Fig. 5c, d; see Methods). CDX Lineage A (L.A) displayed a stable transcriptional signature across the TKI treatment time course (Fig. 4c) and a relative increase in frequency over time (Fig. 4b). CDX Lineages B (L.B) and C (L.C) showed transcriptional rewiring (Fig. 4c) and decreased in frequency after treatment with osimertinib (Fig. 4b). Comparatively, these same three lineages were detectable in vehicle-treated tumors (Fig. 4c) but did not demonstrate changes in abundance over time (Fig. 4b).

Given the stable transcriptional signature of CDX L.A across the time course of the osimertinib treatment and the enrichment of its barcode abundances at osimertinib resistance, we reasoned that through the analysis of single-cell signatures from D0, we could identify resistance-associated transcriptional regulators that pre-exist in treatment-naïve cells. Using iPAGE (see Methods), we found that at D0 CDX L.A shows increased genes from the FOXD1 (FREAC4_01) regulon compared to CDX L.B and L.C (Fig. 5a and Supplementary Fig. 5e, f). The FOXD1 regulon gene set contains genes identified as potential binding sites for the transcription factor, FOXD1, as determined by JASPAR Predicted Transcription Factor Targets^41^, many of which have been independently associated with disease progression, drug resistance, and metastasis in cancer^42–58^.

**Figure 5.**
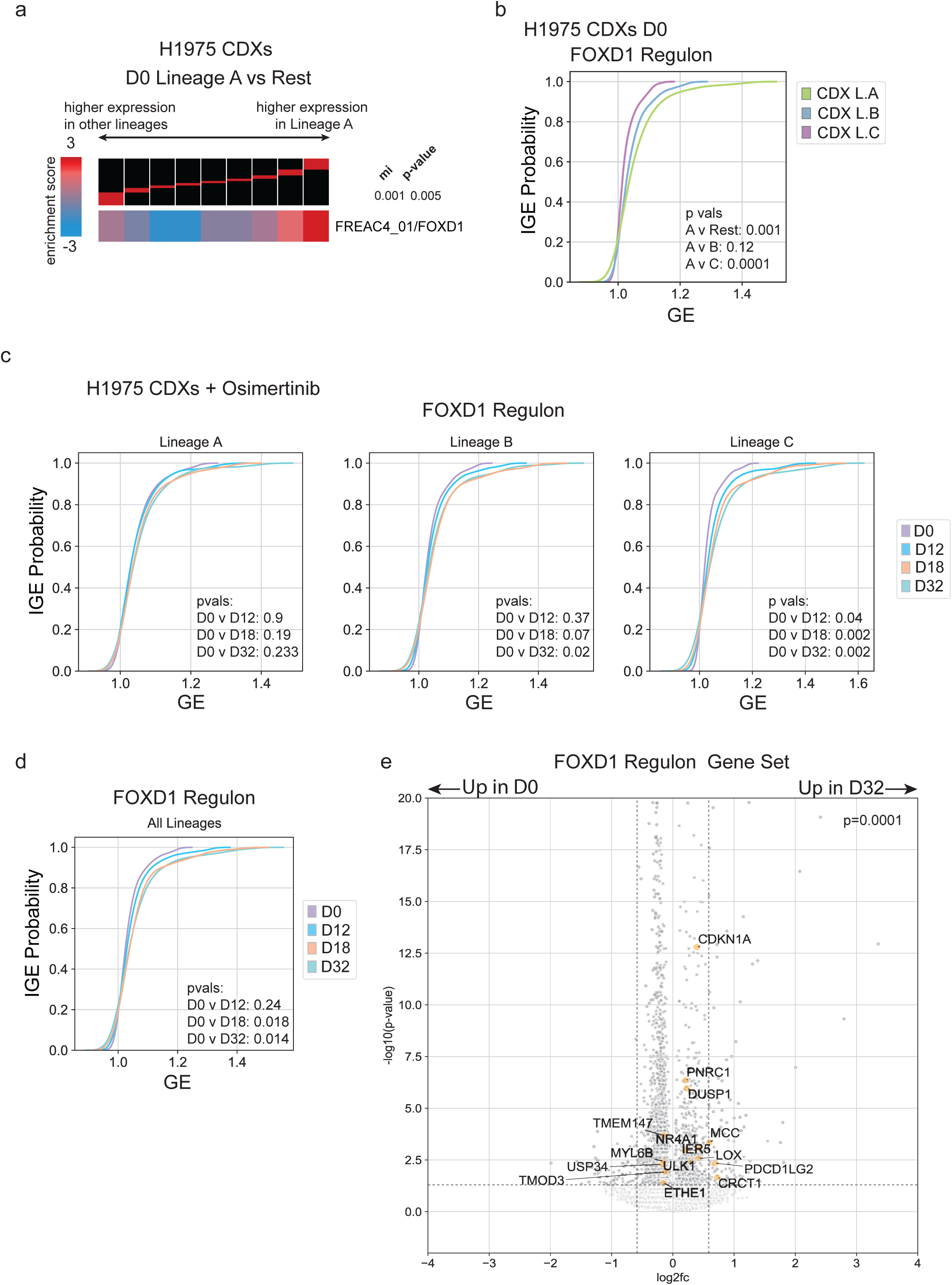
Pathway analysis of gene expression of lineages detected in EGFRmt NSCLC CDXs points to the upregulation of a FOXD1 transcriptional program pre-existing in the lineage population becoming predominant at osimertinib resistance. (a) Pathway analysis of gene expression (iPAGE) comparing CDX Lineage A population versus rest (B-C) at D0 points to the enrichment of FREAC4/FOXD1 regulon. All individual sequenced genes are sorted by log2FC and divided into 9 equally populated bins from lowest (negative) log2FC (left) to greatest log2FC (right). The red bar within the black header indicates the range of log2FC values in each bin. The heatmap is colored by the enrichment or depletion of the gene in each bin. The top/right-most iPAGE bin, containing genes with the greatest log2FC values, is significantly enriched for genes in the FREAC4_01/FOXD1 regulon gene set. The significance of the mutual information value (bits) is calculated using a non-parametric randomization test. (b) Density plots showing significant enrichment of the FREAC4/FOXD1 regulon gene set in CDX Lineage A versus others at D0 (y-axis: IGE Probability, individual gene expression probability, representing the proportion of gene expression values less than or equal to the values in the x-axis; x-axis: GE, gene expression). (c) Density plots for FOXD1 regulon over the course of the osimertinib treatment (D0-D32) show significant enrichment of these modules at D32 CDX Lineages B and C, and stable expression in CDX Lineage A. (d) Density plot for FREAC4/FOXD2 regulon gene set shows significant enrichment of this gene set over time in the entire dataset (y-axis: IGE Probability, individual gene expression probability, representing the proportion of gene expression values less than or equal to the values in the x-axis; x-axis: GE, gene expression). (e) Volcano plot shows differential gene expression between D32 and D0 across all CDX lineages. Yellow annotated genes indicate significantly differentially expressed genes in the FREAC4/FOXD1 gene set. The horizontal dashed line indicates a p-value of 0.05, and the vertical dashed lines indicate +/-log2fc(1.5). (Panels b, c, d, e: p-values calculated using independent t-tests via scipy.stats.ttest_ind). See also Supplementary Figure 5.

Over time, CDX L.A’s expression of the FOXD1 regulon gene set remained stable from treatment-naïve (D0) to treatment persistence (D12) and resistance (D18, D32), while both CDX L.B and L.C significantly upregulated this regulon over treatment time (Fig. 5b, c). This is consistent with the hypothesis that CDX L.A has a relatively stable transcriptional profile during therapy persistence, while CDX L.B and L.C leverage up-regulation of the FOXD1 regulon to adapt to osimertinib treatment, indicating a potential convergent adaptation. Consistent with this hypothesis, we similarly observed significant up-regulation of FOXD1 regulon genes when analyzing all lineages together over time (Fig. 5d, e).

Overall, our lineage tracing approach allowed us to identify a FOXD1 regulon as a significantly upregulated transcriptional program in pretreatment CDX L.A, that also became enriched during the persistent (D12) and resistant (D18, D32) states in the other lineages. This evidence, the reported presence of FOXD1-binding sites in genes associated with disease progression, drug resistance, and metastasis in cancer^42–58^, as well as its role in the proliferation of lung cancer cells and the association with poor patient prognosis in NSCLC^59^, prompted us to perform a functional characterization of CDX L.A and FOXD1 regulon experimentally to confirm whether L.A-enriched CDXs had a survival advantage during osimertinib treatment and whether this advantage pre-existed before therapy in the form of the FOXD1 transcriptional state.

## Enrichment of CDX Lineage A conferred higher progressive disease in xenograft models

The consistent transcriptomic profile of CDX L.A and the increase in relative population frequency across treatment time raised the hypothesis that CDX L.A’s survival advantage pre-existed before treatment in the form of the FOXD1 transcriptional state. We aimed to isolate CDX L.A from bulk barcoded H1975 cells to perform functional experiments upon osimertinib treatment to establish causation in drug persistence and resistance. To do so, we identified surface proteins with high expression in CDX L.A via differential gene expression, and conducted FACS sorting matched with high-throughput sequencing^60^ analyses (Supplementary Fig. 6a, b). Among the surface proteins screened for sorting, GPC4 and SUSD2 had a two-to-four-fold enrichment efficiency in CDX L.A compared to unsorted cells (Supplementary Fig. 6c). We leveraged the higher sorting efficiency of the surface protein SUSD2 to sort the treatment-refractory CDX L.A and performed functional *in vitro* and *in vivo* characterization. Of note, in our prior single-cell RNA sequencing analysis of clinical biopsy specimens, high expression of SUSD2 is also observed in residual disease and therapy persistence in NSCLC patients upon TKI treatment. This makes it a compelling biomarker to characterize disease persistence. Osimertinib-treated CDX L.A-enriched cells (Supplementary Fig. 6d) showed an average three-fold increase in 2D- and 3D-colony number compared to the unsorted cells (U.C.) (Fig. 6a, b and Supplementary Fig. 6d-f), indicating the predicted survival advantage compared to U.C. colonies. We used CDX L.A-enriched cells to generate flank xenografts and treated mice with osimertinib for 26 days until tumors gained maximal size (Fig. 6c). CDX L.A-enriched xenografts were classified at day 26 using RECIST^61^ criteria, which indicated a statistically significant progressive disease phenotype for CDX L.A-enriched compared to unsorted control xenografts (Fig. 6d, e).

**Figure 6.**
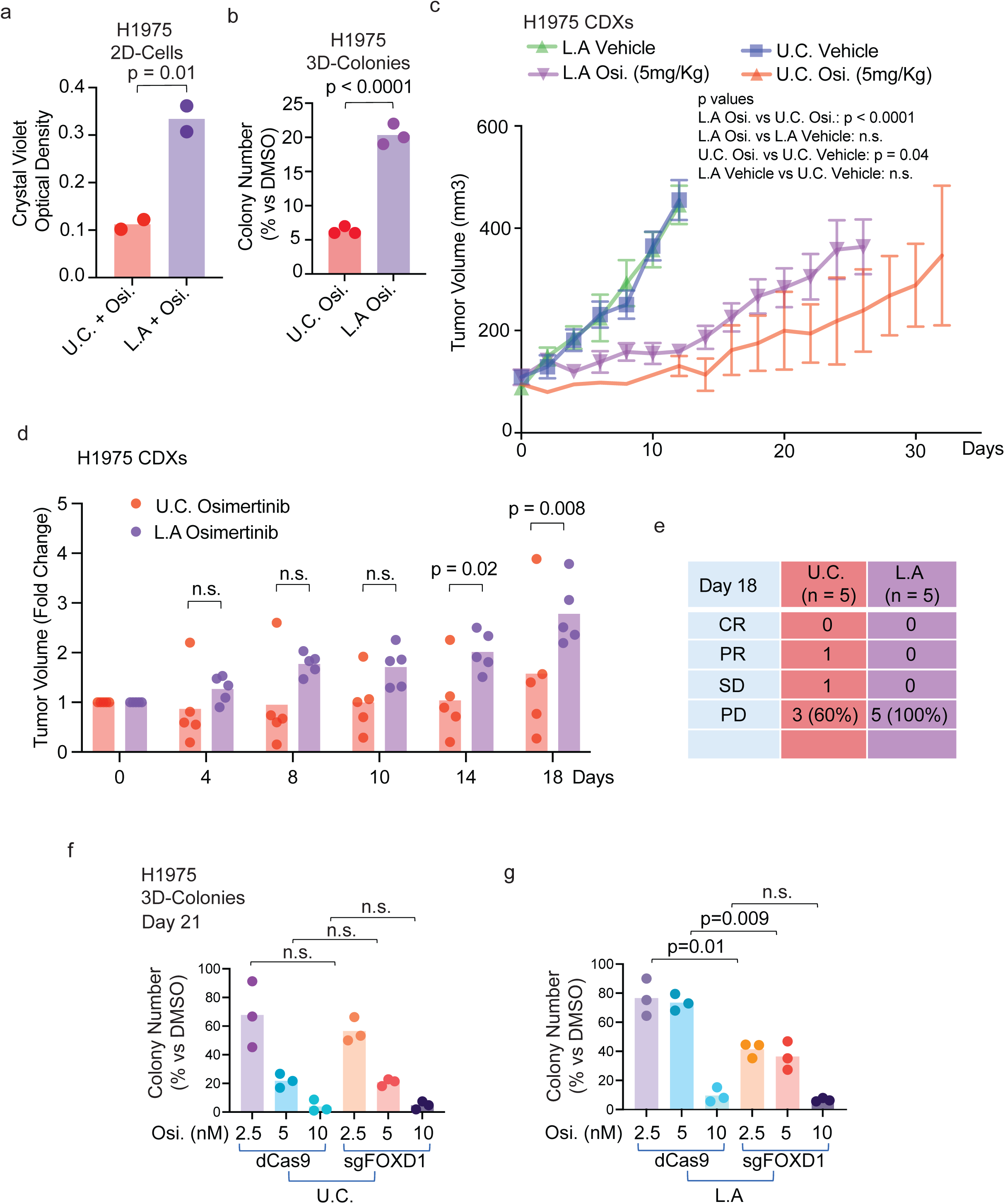
Pre-clinical functional validation of osimertinib-resistant lineages detected in EGFRmt NSCLC CDXs. (a, b) Functional crystal violet assay quantification in 2D- (a) and 3D soft agar colony formation (b) cultures with lineage A enriched (L.A) or unsorted control (U.C.) H1975 cells; quantification of the crystal violet OD (a) or the number of 3D colonies (b) is provided (n = 2-3 replicates per experimental group; p-value calculated using Student’s t-test). (c) Functional in vivo experiments with lineage A enriched (L.A) or unsorted control (U.C.) H1975 CDXs treated with osimertinib or vehicle control (p-value calculated using One-way ANOVA with Tukey’s multiple comparison test). (d, e) The tumor volume fold change was calculated from each xenograft (d) and response to osimertinib classified with RECIST criteria (e) (p-value calculated using Two-way ANOVA with Sidak’s multiple comparison test). (f, g) Functional tests using CRISPR-Cas9 lentiviral infection with sgRNA targeting FOXD1 protein and 3D-soft agar colony formation assay using unsorted control (U.C.) (f) or lineage A enriched (L.A) (g) H1975 cells treated with a dose escalation of osimertinib (Osi., 2.5-10 nM). Colonies were stained with crystal violet at day 21 and quantified (f, g) (n = 3 replicates per experimental group; p-value calculated using Two-way ANOVA with Sidak’s multiple comparison test). (Osi.: osimertinib; CR: complete response; PR: partial response; SD: stable disease; PD: progressive disease). See also Supplementary Figures 6 and 7.

Furthermore, we investigated through whole-exome sequencing whether the L.A population carries or acquires *EGFR* gene alterations compromising therapy response. We did not detect additional *EGFR* mutation or copy number alterations in CDX L.A-enriched xenografts, osimertinib-treated, compared to vehicle-treated controls, nor in CDX L.A-enriched xenografts, osimertinib-treated, compared to osimertinib-treated unsorted control xenografts (Supplementary Table 2), providing evidence against the possibility that acquired or pre-existing modifications of the EGFR gene itself conferred the EGFR TKI resistant phenotype.

In summary, genomic barcoding identified a pre-existing population of clonal cells, CDX Lineage A, with a significant TKI-treatment refractory phenotype in pre-clinical *in vivo* studies. This analysis identified a programmatic biomarker (FOXD1 regulon) of the persistence and enrichment at resistance of CDX L.A during targeted therapy versus the unsorted control xenografts.

## Knockdown of FOXD1 increased sensitivity to osimertinib in NSCLC 3D cell colonies

Next, based on the findings above, we tested the hypothesis that FOXD1 could diminish response to EGFR TKI treatment, specifically in CDX L.A-enriched and unsorted control (U.C.) cells using CRISPRi and a dose-escalation (2.5 - 10 nM) time-course with osimertinib in 3D-soft agar assays (Supplementary Fig. 7a). The efficiency of the sgFOXD1 knockdown was confirmed through qPCR analysis (Supplementary Fig. 7b). Colony growth was monitored at day 7 (Supplementary Fig. 7c), day 14 and day 21 (Supplementary Fig. 7d) and quantified at day 14 (Supplementary Fig. 7e-g) and day 21(Fig. 6f, g and Supplementary Fig. 7g). At day 21 on osimertinib (2.5-5 nM), unsorted control sgFOXD1 cells showed similar degrees of sensitivity to unsorted control dCas9 cells (Fig. 6f). In contrast, CDX L.A-enriched cell colonies in which FOXD1 was silenced had a significant reduction of colony formation compared to L.A dCas9 cells (Fig. 6g). This evidence is in line with the hypothesis of a role of FOXD1 in contributing to resistance in CDX L.A clonal cells.

Our data show that lineages more refractory to osimertinib treatment have a pre-existing engagement of a FOXD1-regulatory program that reduces EGFR inhibitor sensitivity. FOXD1 has been shown to promote cell growth and metastasis in NSCLC^62,63^. A number of the significantly upregulated FOXD1 regulon genes in CDX L.A vs. L.B and L.C at D0 (Supplementary Fig. 5f) and at D32 vs. D0 across all lineages (Fig. 5d) are associated with disease progression, drug resistance, and metastasis in cancer^42–58^. Of these genes, we hypothesize *NR4A1*, which is significantly upregulated at D32 vs D0 across all lineages and significantly upregulated in CDX L.A vs. L.B and L.C at D0 plays a significant role in the FOXD1-related osimertinib resistance identified. NR4A1 is a transcription factor that has been shown to complex with known EGFR-promoter, SP-1, to directly increase transcriptional activity of genes with proximal GC-rich promoter elements such as Survivin and EGFR in lung cancer, pancreatic cancer models, and breast cancer models^64,65^. Consistently, in CDX L.A-enriched cells in which FOXD1 was silenced, 75% of the previously identified FOXD1-regulatory program was downregulated (Supplementary Fig. 8a, b), with the highest significance of downregulation reached for the *NR4A1* gene (Supplementary Fig. 8c). While NR4A1 could contribute to EGFR inhibitor resistance through the mechanisms noted above, future work will be required to elucidate the precise functional role of NR4A1 in EGFR inhibitor resistance.

Next, we confirmed the utility of our lineage tracing scRNA-seq approach in identifying FOXD1 and SUSD2 as markers of pre-existing drug-resistant lineages by comparing our barcode-derived transcriptional clusters to standard practice clustering of single-cell RNAseq based on gene expression alone^37^. Clustering of the single-cell RNAseq gene expression CDX data identified thirteen clusters (Supplementary Fig. 9a). Clusters 5 and 9 were identified as dying cells and omitted from further analysis. No cluster increased in relative population frequency over the entire treatment time. However, Clusters 0, 6, and 8 had a net increase in population frequency over time (Supplementary Fig. 9a). Additionally, when the genes from the top FREAC4/FOXD1 Regulon iPAGE bin (Supplementary Fig. 5f) and SUSD2 (Supplementary Figure 6a) were plotted across clusters, no single cluster fit the CDX L.A transcriptomic profile (Supplementary Fig. 9b, c).

The data indicate that our genomic-barcode-based lineage tracing technique was essential to identify CDX L.A and, subsequently, FOXD1 as a potential marker and promoter of TKI resistance.

## Clinical patterns of Hh signaling and FOXD1 regulon expression validate preclinical findings

To confirm the clinical relevance of the Hh pathway and FOXD1 programs identified in our osimertinib-resistant lineages before treatment in our PDO and CDX datasets, we examined EGFRmt NSCLC-specific single-cell RNA-sequencing clinical data from our group^5^. In EGFRmt NSCLC primary tumors, expression of the Hallmark Hedgehog Signaling gene set is higher in treatment-naïve (TN) human tumors compared to tumors at residual disease (RD) after TKI treatment, aligning with the PDO L.D expression pattern (Fig. 7a). This evidence suggests that the Hh pathway upregulation pretreatment might indicate the presence of a plastic state that primes cells to adapt after TKI therapy transcriptionally. Furthermore, RD tumor samples significantly upregulate *SNAI1* and show trends toward *VIM* upregulation (Fig. 7b, c). SNAI1 is a positive regulator of EMT^66^. VIM is a known marker of cells that have undergone EMT^67^, suggesting these cells undergo a transcriptional rewiring, reflecting the signals observed in the PDO L.D. expression pattern.

**Figure 7.**
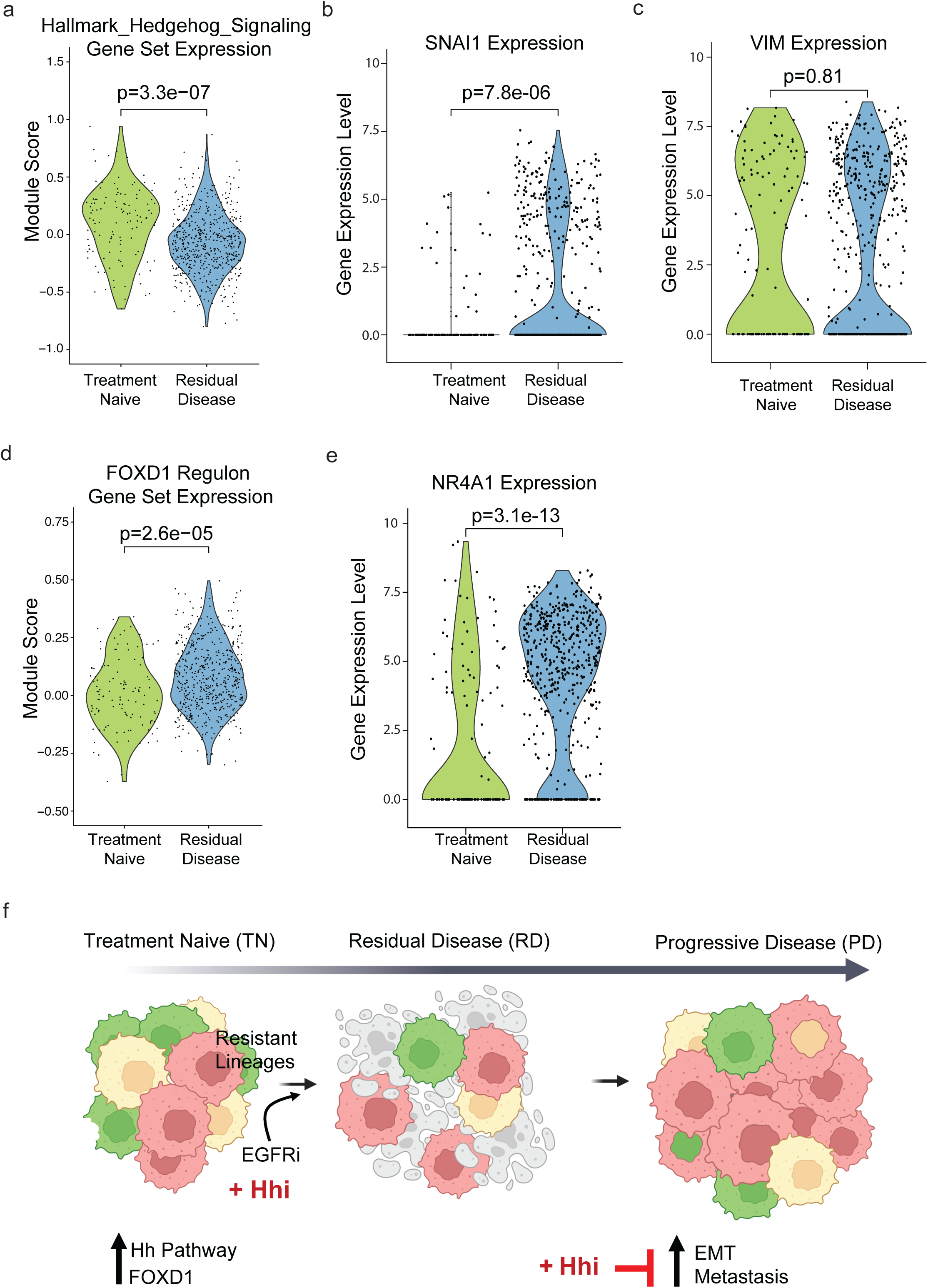
Cross-analysis of osimertinib-resistant lineage’s signatures with clinical datasets. (a) Hedgehog signaling module’s representation in Smart-Seq2 scRNAseq dataset from internal UCSF cases treated with EGFR targeted therapy, at treatment naïve or residual disease states (p-values calculated with Welch Two Sample t-test). (b-c) Gene expression level for EMT biomarkers *SNAI1* and *VIM* in Smart-Seq2 scRNAseq dataset from internal UCSF cases treated with EGFR targeted therapy, at treatment naïve or residual disease states (p-values calculated with Welch Two Sample t-test). (d) FOXD1 regulon representation in Smart-Seq2 scRNAseq dataset from internal UCSF cases treated with EGFR targeted therapy, at treatment naïve or residual disease states (p-values calculated with Welch Two Sample t-test). (e) Gene expression level for FOXD1 gene target *NR4A1* in Smart-Seq2 scRNAseq dataset from internal UCSF cases treated with EGFR targeted therapy, at treatment naïve or residual disease states (p-values calculated with Welch Two Sample t-test). (f) Model diagram describing osimertinib-resistant lineages identified in our studies, characterized by an a priori up-regulation of Hedgehog (Hh) pathway biomarkers and FOXD1 regulon in treatment naïve EGFRmt NSCLC tumors, leading to therapy resistance and tumor progression (EMT, ECM-metastasis). A combination of Hhi and osimertinib resulted in more effective suppression of EMT transition and durable responses in primary organoids.

Within EGFRmt NSCLC patients, the FOXD1 regulon and our top candidate gene, *NR4A1*, were significantly enriched at the drug-tolerant RD tumor state compared to treatment-naive tumors (Fig. 7d, e). These clinical specimen data align with the pattern shown by the CDX model that this lineage is less sensitive to EGFR inhibitor treatment and that all lineages upregulate the FOXD1 regulon in response to treatment. These data support the potential contribution of the FOXD1 regulon to therapy persistence and eventual resistance in a clinical setting.

### Discussion

Single-cell RNA sequencing approaches applied to clinical specimens have broadened our understanding of tumors’ compositional and state heterogeneity in response to treatment. However, the limited cell number derived from each patient’s biopsy and the lack of information on the evolution of clonal populations upon therapy in different model systems limited the characterization of the heterogeneous transcriptional programs of tumor cells in the TN, RD, and PD states.

We overcame these limitations using our static barcoding approach matched with single-cell RNAseq on highly heterogeneous primary organoid NSCLC samples and in vivo models to track clonal evolutionary trajectories and reveal otherwise occult distinct seeds of pre-existing, TKI-treatment-refractory lineages that are selected over the course of the TKI treatment (Fig. 7f). Through analysis of TN, RD, and PD states with increased granularity, we were able to identify and study treatment-refractory clones and mechanisms of tumor cell adaptation leading to therapy resistance. Our GBC lineage tracing approach uncovered the presence of lineages in TN tumors with reduced sensitivity to therapy (Fig. 7f). To our knowledge, this is among the first reports on using single-cell lineage tracing matched with single-cell RNA sequencing approaches in primary and *in vivo* NSCLC models during therapy to inform actionable programs to delay the acquisition of therapy resistance. By identifying the transcriptional programs of these resistant lineages before treatment, we were able to develop pharmacologic combinatorial treatments and genetic approaches to intercept adaptive programs to reduce the reservoir of drug-persistent tumor cells and induce more durable responses.

Specifically, we identified a subset of treatment-refractory lineages with Hh-pathway and FOXD1-regulon-dependent phenotypes, for which we provided functional characterization through pharmacological and genetic inhibition. Single-cell RNA sequencing analyses of EGFRmt NSCLC clinical cases in the treatment-naïve and residual disease subgroups confirmed a bimodal regulation of the Hedgehog regulon and EMT biomarkers, *SNAI1* and *VIM*, with Hh regulon being significantly decreased and EMT biomarkers enriched at the RD state compared to TN cells (Fig. 7a-c), in line with our preclinical data. We show that combining Hh inhibitor with osimertinib can enhance the response to EGFR inhibitor therapy in primary EGFRmt NSCLC organoids. Based on the bimodal pathway data, we propose this treatment effect is due to the ability to target this Hh positive subpopulation of cells that are primed for transcriptional plasticity and persistence and more potently suppress the adaptation toward the subsequent therapy-persistent EMT phenotype (Fig. 3). We identified FOXD1 regulon and its target gene *NR4A1* as potential biomarkers of osimertinib RD in single-cell RNA sequencing analyses of EGFRmt NSCLC cases (Fig. 7d, e), in line with our evidence of persistence and enrichment of FOXD1 regulon expression over the course of the osimertinib treatment in our identified *in vivo* CDX lineage populations. Identifying and targeting transcriptional and cell state plasticity and priming events and programs are essential goals in cancer biology and therapeutics research, which our findings may help to address.

Overall, identifying transcriptional programs linked to treatment-refractory populations as we have executed here using orthogonal high-resolution approaches in preclinical and clinical systems has the potential to enable better prediction of successful combinatorial treatments for cancer patients for clinical translation.

### Online Methods

## Cell lines, reagents, organoid cultures, and in vivo cell-derived xenografts

All cell lines were obtained from the American Type Culture Collection (ATCC), authenticated through STR profiling, and confirmed negative for mycoplasma contamination. Cell lines were cultured in a humidified incubator with 5% CO2 at 37 °C. HEK293-FT cells were cultured in DMEM (HyClone, GE Healthcare) supplemented with 10% FBS, 0.1X penicillin, and streptomycin. Primary, patient-derived EGFRdel19 positive organoid cells TH-107 were derived from a tumor biopsy and cultured in matrigel as previously described^68^. The EGFR L858R, T790M H1975 cells were grown in RPMI 1640 medium (HyClone, GE Healthcare), containing 10% FBS (SAFC, Sigma-Aldrich) and 1X penicillin and streptomycin (HyClone, GE Healthcare) per datasheet recommendations. Osimertinib and sonidegib were purchased from Selleckchem. Immunoblotting antibodies were EGFR (Cell Signaling Tech., 4267), EGFR E746-A750del/Del19 (Cell Signaling Tech., 2085), EGFR L858R (Cell Signaling Tech., 3197), pEGFR-Tyr1086 (Thermo Fisher Sci., 369700), p-EGFR-Tyr1068. (Cell Signaling Tech., 3777), p-ERK (Cell Signaling Tech., 4370), p-Akt (Cell Signaling Tech., 4060), ERK (Cell Signaling Tech., 9102), Akt (Cell Signaling Tech., 2920), cleaved PARP (Cell Signaling Tech., 9541), Smoothened (Cell Signaling Tech., 92981), Vimentin (Cell Signaling Tech., 3932), E-Cadherin (Cell Signaling Tech., 3195), Claudin-1 (Cell Signaling Tech., 4933), Slug (Cell Signaling Tech., 9585), Actin (Sigma, A2228). All antibodies were diluted per datasheet recommendations.

Barcoded TH107 organoid cells were treated with osimertinib 500 nM for 37 days, generating osimertinib persisters and collecting single-cell suspensions at days 0 (T0), 7 (T1), 14 (T2), 21 (T3), and 30 (T4) for analysis. For the 3D-CTG assay, barcoded TH107 organoid cells were treated with a dose escalation of osimertinib (1 - 10000 nM), sonidegib (1 - 10000 nM), or a combination for 5 days. For the long-term treatment, barcoded TH107 organoid cells were treated with a dose escalation of osimertinib (1 - 100 nM), sonidegib (10 uM), or a combination for 21 days, replenishing the drugs once per week, after which organoids were harvested, counted, and pellets used for protein extractions and immunoblotting.

Barcoded H1975 xenografts were established following UCSF IACUC IRB-approved protocols. H1975 flank xenografts were generated by injecting 1 million cells into the flank of 4-week-old female athymic Nude mice. After mice randomization, osimertinib was administered daily by oral gavage at 5 mg/Kg. The xenografts were measured using a digitized caliper, and mice were euthanized when tumors reached a diameter of 20 mm. Vehicle-treated control CDXs were collected on day 12 after reaching their maximum size. Unsorted control tumors were resected at four time points along the course of the TKI treatment to derive single cells: at day 0 (T0), 12 (T1), 18 (T2), and 32 (T3) (Fig. 1e).

## Transfection optimization and lentiviral barcoding of NSCLC primary and patient-derived cells

The barcoding lentiviral construct library, containing approximately 100,000 unique, 18-nucleotide static genomic barcodes, was a gift from Weissman’s lab^69^. HEK293-FT cells were transfected with the vectors using Mirus reagent, following the datasheet recommendations. After 72 hrs of transfection, the virus was collected and used to transduce NSCLC cells in a medium containing polybrene (8 μg/ml; Sigma-Aldrich). Selection of successfully transduced cells was carried out using puromycin (1 μg/mL; Gibco) or sorting for BFP-positive cells for the barcoded cells.

We optimized the barcoding efficiency in our NSCLC preclinical model’s genomes, titrating the lentiviral transducing library collected from the HEK293 cells’ transfection. Five million TH107 PDO cells were incubated with different percentages of viral-containing media, respectively, 25%, 50%, and 100%. The 50% condition resulted in high PDO transduction efficiency with less toxicity, as assessed by microscopic observation of BFP-positive cells (Supplementary Fig. 1b). FACS-sorted BFP-positive, barcoded H1975 cells (Supplementary Fig. 1e) were seeded at different densities, ranging from 500 cells/well to 64,000 cells/well, transduced, and sequenced via MiSeq^60^. Miseq’s results indicated that beginning with a 2,000 founder population of H1975 cells leads to the recovery of 619 unique barcodes (Supplementary Fig. 1f). In both PDO and H1975 cells used to derive xenograft models, we proceeded with ∼2000 and ∼1000 uniquely barcoded clonal populations (lineages), respectively, to balance the diversity of the lineage pool against the sampling limitations of single-cell RNA sequencing, and banking early passages cells.

## Preparation of the libraries for sample pooling, gDNA, and scRNA-seq analyses

Single-cell suspensions collected from our in vitro PDO and in vivo CDX models were divided, with 10% of cells used for single-cell RNA (scRNA-seq) and the remaining 90% of cells used for genomic bulk sequencing (Fig. 1a). Prioritization of the samples for gDNA amplicon sequencing after scRNA-seq allowed the genomic barcode (GBC) frequencies, calculated more accurately from the genomic bulk sequencing at each time point, to be mapped back to the transcriptional information from the scRNA-seq. Two additional single-cell libraries were made to amplify the GBCs from the 10X scRNA-seq library and to amplify the Multiseq tags^70^ used for sample hashing. Integration of bulk GBC frequencies and single-cell transcriptomic data allowed downstream analysis to assess how the osimertinib treatment and evolution of lineage-specific transcriptional signatures modulated lineage representation and abundances over time.

## Genomic barcode (GBC) cleaning

Inherent to scRNAseq-based methods of lineage capture is the low number of cells that can be sequenced from the sample compared to genomic sequencing-based methods. Estimating lineage-based population frequency from counting single cells limits the number of lineages that can be effectively traced through time and requires a high degree of sequencing and single-cell investment^71^. For these reasons, we added paired gDNA to lineage tracing-based scRNA-seq data collection and performed PCR-based targeted enrichment of lineage tracing barcodes. This allowed for estimating population frequencies among tens of millions of cells by counting the lineage barcodes directly. The gDNA barcodes are then enriched in the scRNA-seq fraction by a similar PCR-based targeted enrichment from cDNA. Because the barcodes are proximal to the 3’ end of the transcript on an introduced transgene, it allowed for selective amplification and seamless compatibility with the 10X 3’ capture-based system. This way, we could confidently proceed with hundreds of unique lineages and obtain high accuracy in measuring population frequency while using a relatively low number of cells for single-cell RNA sequencing. After expansion, we froze large and early banks of these models to allow for genetic drift.

We recovered 72,319 unique GBCs from TH107 PDO across all samples. GBC frequencies were calculated for each sample individually. To create a clean set of GBCs for integration with transcriptional data, we began with the 34,889 unique barcodes recovered from the T0 PDO sample. We then filtered for barcodes that occurred at least twice at T0 and were present at all other time points, leaving 2,384 barcodes to be overlaid onto the scRNA-seq data (Supplementary Fig. 2a). We recovered 1,035,878 cells from H1975 CDXs across all samples. After ranking barcodes by count frequency and binning them according to decreasing counts, the appropriate bin size was determined by setting an FDR of 0.05% for each sample, returning 10,054 unique barcodes. After finding the intersection of shared barcodes between all CDX samples, we recovered a list of 1096 genomic barcodes for downstream processing.

## Sample label integration and single-cell object generation

Sample-specific hashtag oligos (HTOs) were extracted with corresponding single-cell 10X cell barcodes, and sample assignments were called using the scEasyMode pymulti package. Lineage tracing genomic barcode files were split into 16 smaller files for faster processing, and 10X cell barcode, 10X UMI, and lineage tracing barcode were extracted for each read. Reads were filtered by comparing sequenced 10X cell barcodes and known 10X cell barcodes and by filtering for lineage tracing GBCs with a count greater than 400; scEasyMode pymulti was run to call appropriate lineage tracing barcodes. Bulk lineage tracing barcode frequencies were calculated by dividing individual barcode count values by the sum of all lineage tracing barcode counts. Lineage tracing barcodes and sample identities were then integrated into the single-cell object based on matched 10X cell barcodes. Lineage tracing barcode frequencies calculated from bulk sequencing were overlaid onto the single-cell object. The annotated single-cell object was then filtered for cells with >90% human genes, <25% mitochondrial genes, and <2500 gene counts.

## Lineage cluster identification

Genomic Barcode (GBC) frequencies were calculated as a percent of the total GBC count per timepoint for each treatment arm and normalized by dividing each GBC frequency in osimertinib-treated samples by the frequency in DMSO-treated samples. For identifying clusters in barcoded TH107 PDO cells, each GBC, relative frequency scores were calculated by dividing the frequency of osimertinib-treated cells by DMSO-treated cells on day 7 (T0) and day 14 (T2). These scores were hierarchically clustered to group cells with similar expansion patterns to identify PDO lineages A, B, C, and D. For identifying clusters in barcoded H1975 CDX cells, GBC frequencies were calculated from total bulk GBC sequencing. The occupancy score was calculated for GBC_Cluster by finding the proportion of each GBC in each Leiden cluster at each time point. GBC_Cluster values were hierarchically clustered to identify CDX Lineage groups A, B, and C. These values were visualized by PCA and colored by lineage identity.

## Identifying differentially regulated gene sets

For CDXs and PDOs, differentially expressed genes defining each lineage at T0, before osimertinib pressure, were extracted via scanpy.rank_gene_groups. The log2 fold-change (log2fc) values and gene names were fed into iPAGE^24^, and all sequenced genes were sorted and binned according to their log2fc values. Briefly, iPAGE leveraged mutual information scores to identify significantly informative pathways and hypergeometric distribution to determine significant levels of over- or under-representation of each pathway within a log2fc bin^24^.

## Soft agar, crystal violet, and 3D-CTG assays

Soft agar colony formation assay was performed as described before^72^. The colony number was counted after 2-3 weeks of growth using ImageJ software (NIH) after incubating wells with crystal violet 0.005% in water and acquiring plate scans (ImageQuant, GE Healthcare). For the 3-day 2-D crystal violet assay, we applied a 0.1% water crystal violet stain solution after drying and fixing cells with PFA 4%. The plates were then scanned using ImageQuant (GE Healthcare), and the crystal violet was quantified by spectrophotometer after dissolving the cell-bound-crystal violet in a 95% ethanol solution. 3D-CTG (Promega) was used to assess osimertinib and sonidegib IC50 in TH107 organoid cells, following the datasheet recommendations. Long-term (three weeks) functional studies using barcoded TH107 treated with osimertinib and sonidegib were performed as described above.

## CRISPR/Cas9 assays, Fluorescent-Activated Cell Sorting (FACS), qPCR analyses, and immunoblotting

Following the lentiviral barcoding protocol described above, we transduced barcoded H1975 cells with CRISPRi constructs and sgRNAs for FOXD1. We sorted for mCherry positive cells for the CRISPR/dCas9 positive cells or double BFP positive and mCherry positive cells for the sgRNAs positive and CRISPR/dCas9 positive cells. The effective CRISPRi knockdown was confirmed by qPCR assay.

Fluorescent-activated cell sorting to purify transduced cells was performed as previously described using a FACS Aria-II cell sorter^73^. Briefly, cells were trypsinized, resuspended in 2% FBS in PBS FACS buffer, filtered through a 45 μm strainer, and then sorted.

qPCR analysis was carried out on cDNAs synthesized with SensiFast cDNA Synthesis Kit (Bioline) following the manufacturer’s recommendations, using 1 mg RNA purified from the transduced cells (RNeasy Mini Kit, Qiagen). A 1:3 cDNA dilution was used for each qPCR reaction. Four replicates per sample were used in the qPCR assay using SYBR-Green chemistry and primers purchased from IDT. The gene expression levels were quantified with the QuantStudio 12K Flex Software (Applied Biosystems), and data are presented with the 2^ ^ΔΔCt^ method with endogenous GAPDH control. Immunoblotting analysis was carried out as previously described^74^.

## RNA-seq, Data Processing, and Differential expression analysis

Bulk RNAs were extracted from H1975 L.A-enriched cells carrying sgFOXD1 or dCas9 control constructs (Qiagen RNeasy kit) and sequenced by Novogene (NovaSeq X Plus Series, PE150). After raw reads alignment using the HISAT2 program^75^, gene expression levels were estimated by the transcript counts that mapped to the genome or exon, and Fragments Per Kilobase of Transcript sequence per Million base pairs sequenced (FPKM) were computed^76^. Differential gene expression analysis (DESeq2 package, Bioconductor) was used in R (v.4x), considering the gene’s expression change significant when adjusted p-value < 0.05. Data visualization included volcano plots, bar plots, dot plots, and heatmaps using ggplot2, pheatmap, ggrepel and EnhancedVolcano packages with gene annotations incorporated using biomaRt and org.Hs.eg.db packages.

## Investigating clinical relevance

Single genes relevant to each gene set were selected and visualized via a KM-plot^77^. Samples were subset to include only adenocarcinoma lung patients, and the follow-up threshold was set to 120 months. The overall survival cohort included 1161 patients. The progression-free survival cohort included 906 patients.

The expression of complete gene sets of interest was investigated by subsetting EGFRmt NSCLC lung tumor cells from the scRNAseq data set from previous work^5^. Module scores were calculated via Seurat’s^78^ AddModuleScore function, and statistics were calculated by Welch Two Sample t-test.

## Statistical Tools

Statistical significance for the functional tests was calculated using Two-way ANOVA with Sidak’s multiple comparison test, one-way ANOVA with Tukey’s multiple comparison test, or Student T-test (GraphPad Prism vs 10.3.0). All statistics for the biocomputational analyses are provided in the text and legends to the figures.

## Acknowledgments

HG is an Arc Core Investigator, and research in their lab is supported by the Arc Institute. This study was supported in part by the Arc Institute. We thank Brian Plosky and Chiara Ricci-Tam from Arc for the helpful feedback on the manuscript.

