## Supplemental Figures 1-9 for "Actionable biological programs to enhance EGFR-targeted therapy response unveiled by single-cell lineage tracing in clinically relevant lung cancer models"

Supplementary Figures 1-9

Supplementary Figure 1

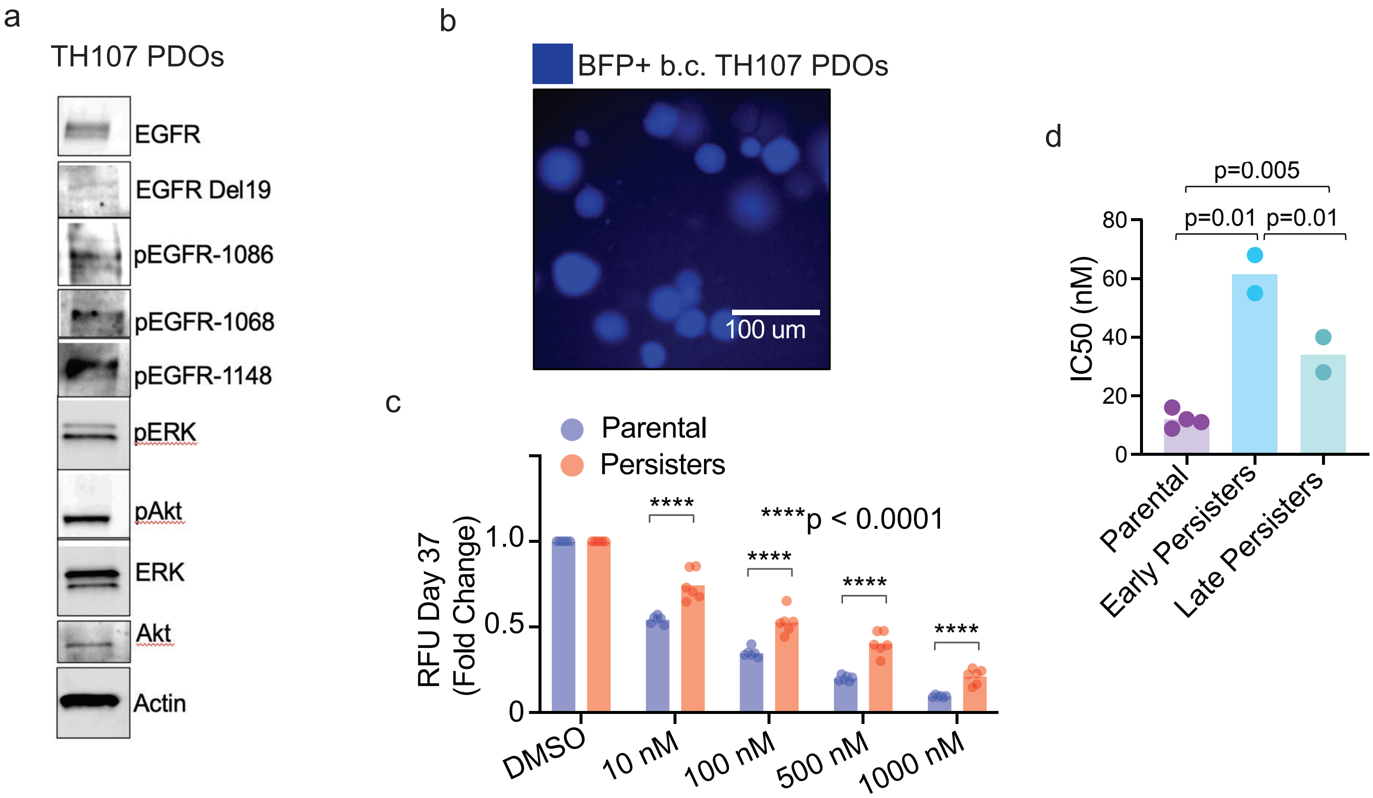

**Supplementary Figure 1**

(a) Biochemical analysis of EGFR signaling in TH107 PDOs. (b) Immunofluorescence for Blue Fluorescent Protein (BFP) reporter gene attesting successful expression of the barcoding construct in barcoded (b.c.) TH107 PDOs. (c) Parental and persisters (at day 37 on treatment) TH107 PDOs quantification, using 3D-CTG and a dose escalation of osimertinib (10 nM - 1000 nM) (p-value calculated using Two-way ANOVA with Sidak's multiple comparison test). (d) IC50 values of TH107 PDOs on osimertinib (500 nM), collected at D14 and D30 (p-value calculated using one-way Anova and Tukey's multiple comparisons test).

Supplementay Figure 2

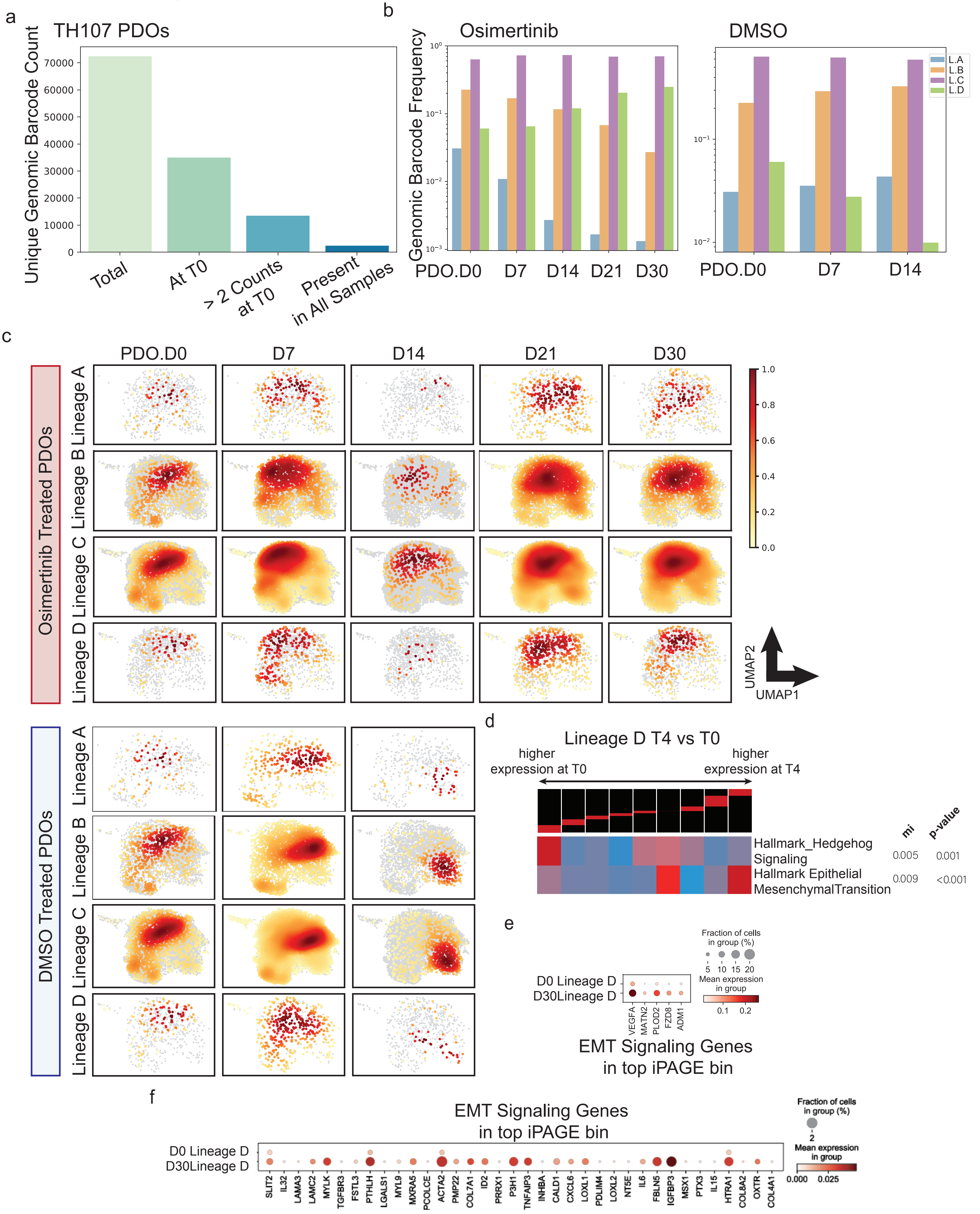

**Supplementary Figure 2**

(a) The filtering of unique genomic barcode count from TH107 PDOs. (b) Genomic barcode frequency scores for L.A-L.D populations highlight a gradual enrichment of the L.D population upon osimertinib treatment compared to DMSO control. (c) UMAP projections of lineage population A, B, C, and D in osimertinib (top) and DMSO (bottom) treatment groups over the course of the in vitro TH107 PDO experiment (PDO.D0-D30), showing distinct transcriptional phenotypes. (d) iPAGE plot comparing the L.D population at D30 vs the L.D population at D0. The bottom/leftmost iPAGE bin is enriched for genes in the Hallmark Hedgehog Signaling gene set; the top/right-most iPAGE bin is enriched for Hallmark EMT genes. (e, f) Dotplots of EMT Signaling Gene Set genes present in the top iPAGE bin. (Panel d: significance calculated using non-parametric randomization – see Methods).

Supplementary Figure 3

a

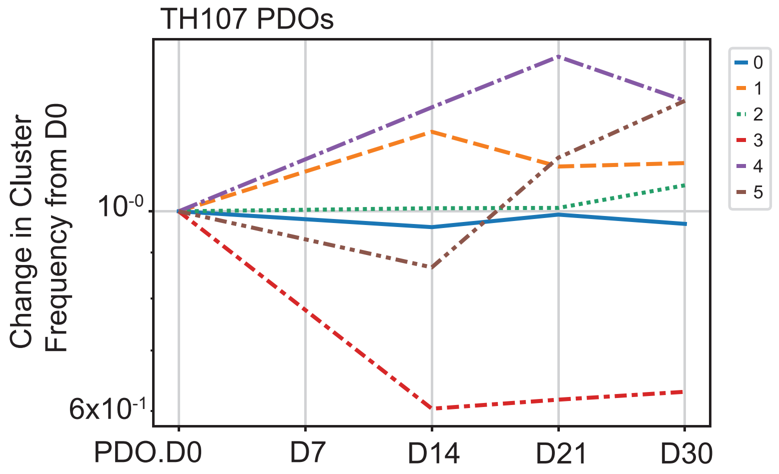


b

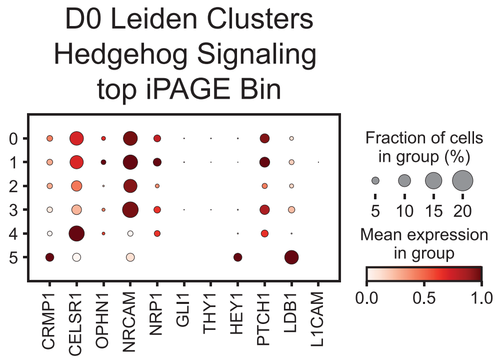


c

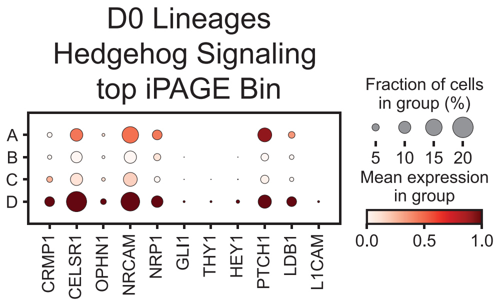

**Supplementary Figure 3**

(a) Leiden clustering of the TH107 PDO data identified six clusters (clusters 0-5). (b, c) Dot plots with single-cell RNAseq data from TH107 PDOs obtained by checking the expression of all the Hallmark Hedgehog Signaling genes from the top iPAGE bin in each of the six Leiden clusters (b) and in lineage A-D populations identified by lineage tracing (c).

Supplementary Figure 4

a

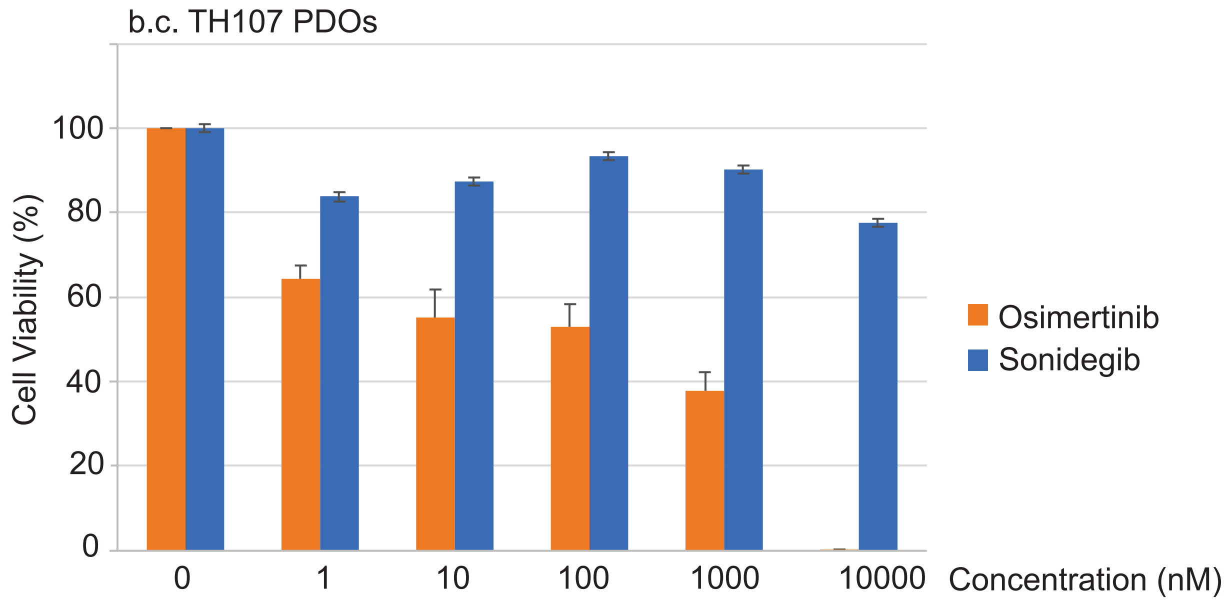


b

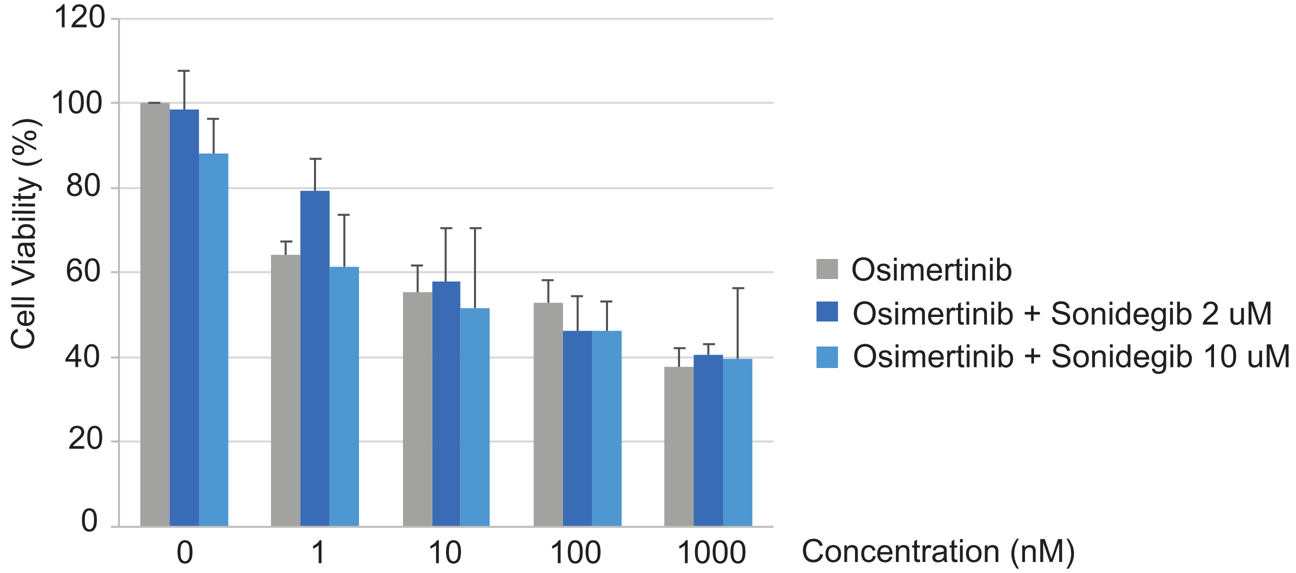


c

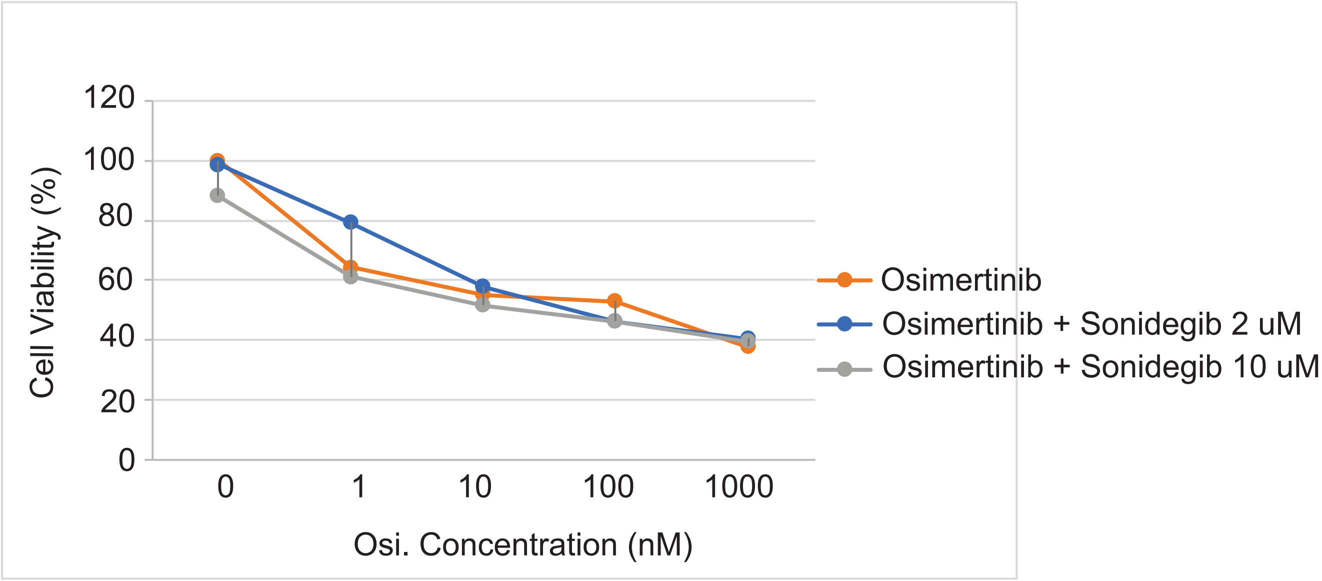

**Supplementary Figure 4**

(a-c) Pharmacological inhibition of Hedgehog pathway using sonidegib in combination with osimertinib in acute treatment (5 days) 3D-CTG with barcoded TH107 PDOs (error bars representing the standard deviation).

Supplementary Figure 5

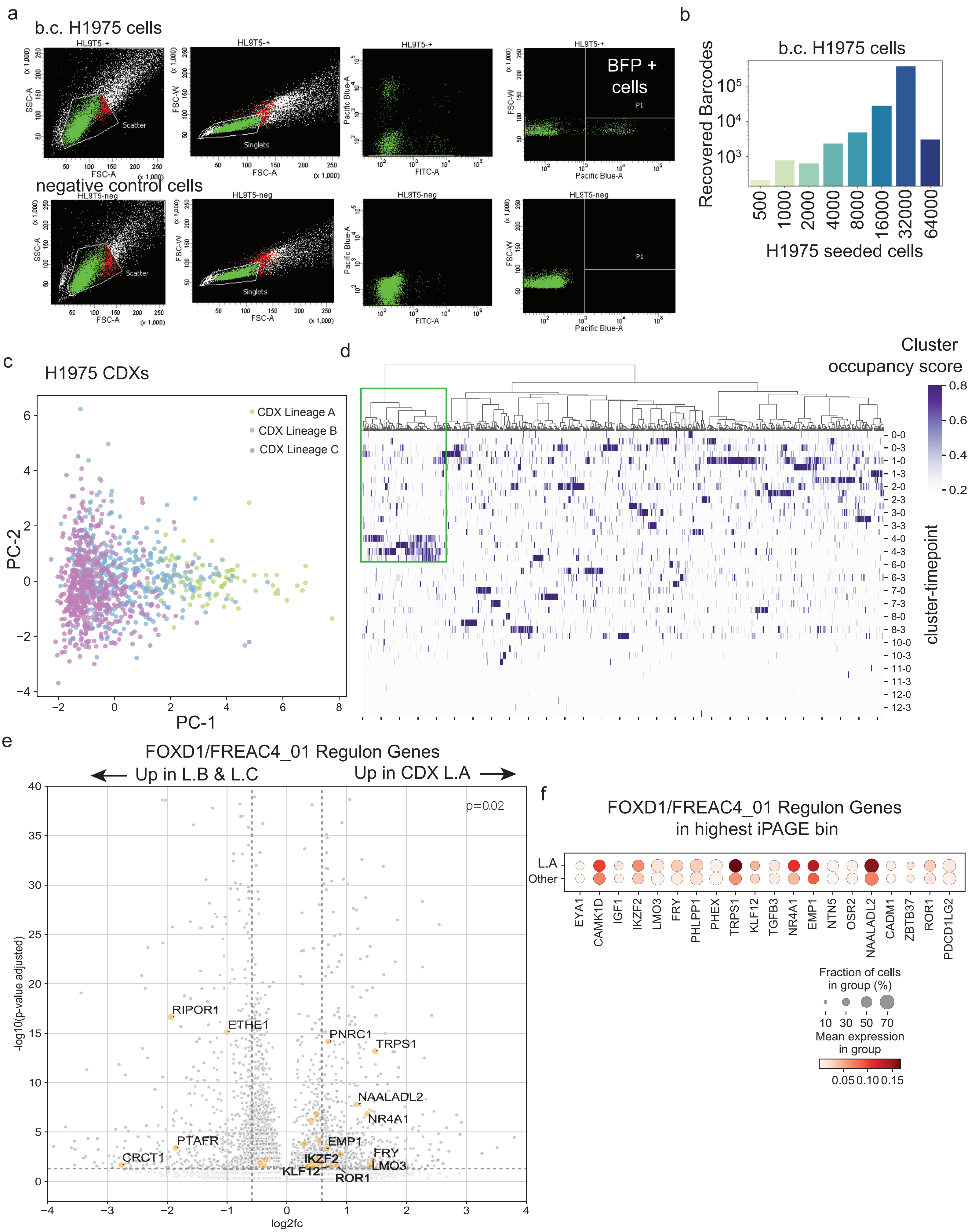

**Supplementary Figure 5**

(a) Sorting of BFP + cells from barcoded (b.c.) H1975 cells transduced with the lentiviral barcoding system. (b) Recovered barcodes from H1975 cells transduced with the lentiviral barcoding system using a serial dilution of seeded cells. (c-d) Principal component analysis (c) and identification of three distinct lineage populations (L.A, L.B, L.C) with different gene expression clusters (d) in H1975 CDXs (lineage A population indicated in the green area in d). (e) Volcano plots showing enrichment of FREAC4/FOXD1 regulon gene set in L.A population at D0 from H1975 CDXs. Yellow dots represent significant FREAC4/FOXD1 regulon genes. Vertical dashed lines represent +/- log2(1.5)FC. The horizontal dashed line represents the log10(0.05) p-value. FOXD1 regulon genes with FC > 1.5 and p-value < 0.05 are annotated (p-values calculated using independent t-tests via scipy.stats.ttest_ind). (f) FREAC4/FOXD1 regulon gene set belonging to the top iPAGE expression gene bin enriched in lineage A (L.A) population from H1975 CDXs.

Supplementary Figure 6

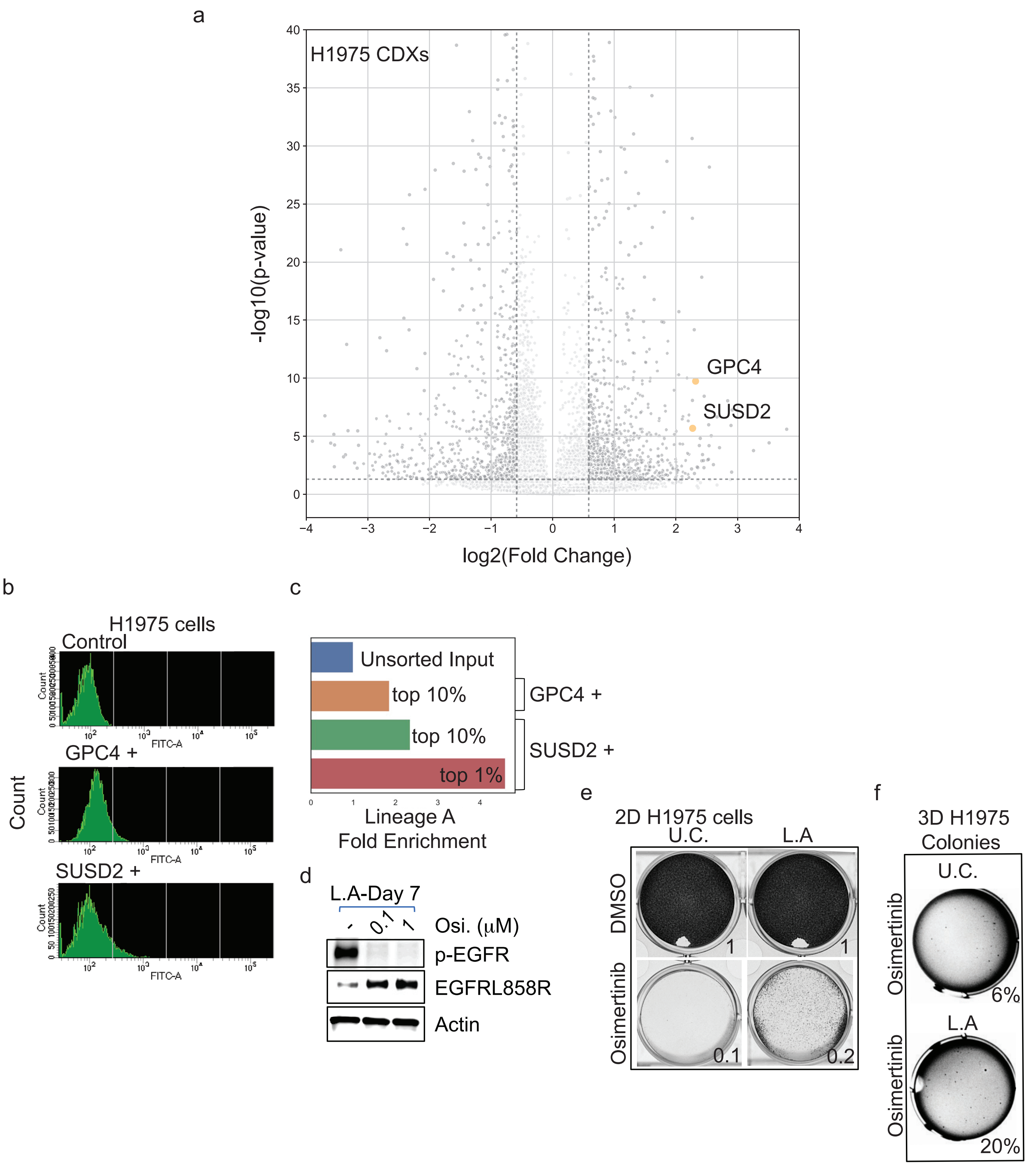

**Supplementary Figure 6**

(a) Volcano plots depicting statistically significant enrichment of surface proteins GPC4 and SUSD2 in lineage A population from H1975 CDXs (p-values calculated using Scanpy's default T-test overestimated variance). (b, c) Sorting SUSD2 + or GPC4 + cells from barcoded H1975 cells (b) and enrichment score of lineage A barcodes post-sorting (c). (d) Biochemical analysis attesting p-EGFR inhibition upon treatment with Osimertinib (Osi.) in lineage A enriched (L.A) H1975 cells. (e, f) Representative images of functional crystal violet assays in 2D- (e) and 3-D soft agar colony formation (f) cultures with lineage A enriched (L.A) or unsorted control (U.C.) H1975 cells; average quantification of the crystal violet OD or the percentages of 3D osimertinib-treated colonies vs. DMSO-treated colonies is provided (n = 2-3 replicates per experimental group).

Supplementary Figure 7

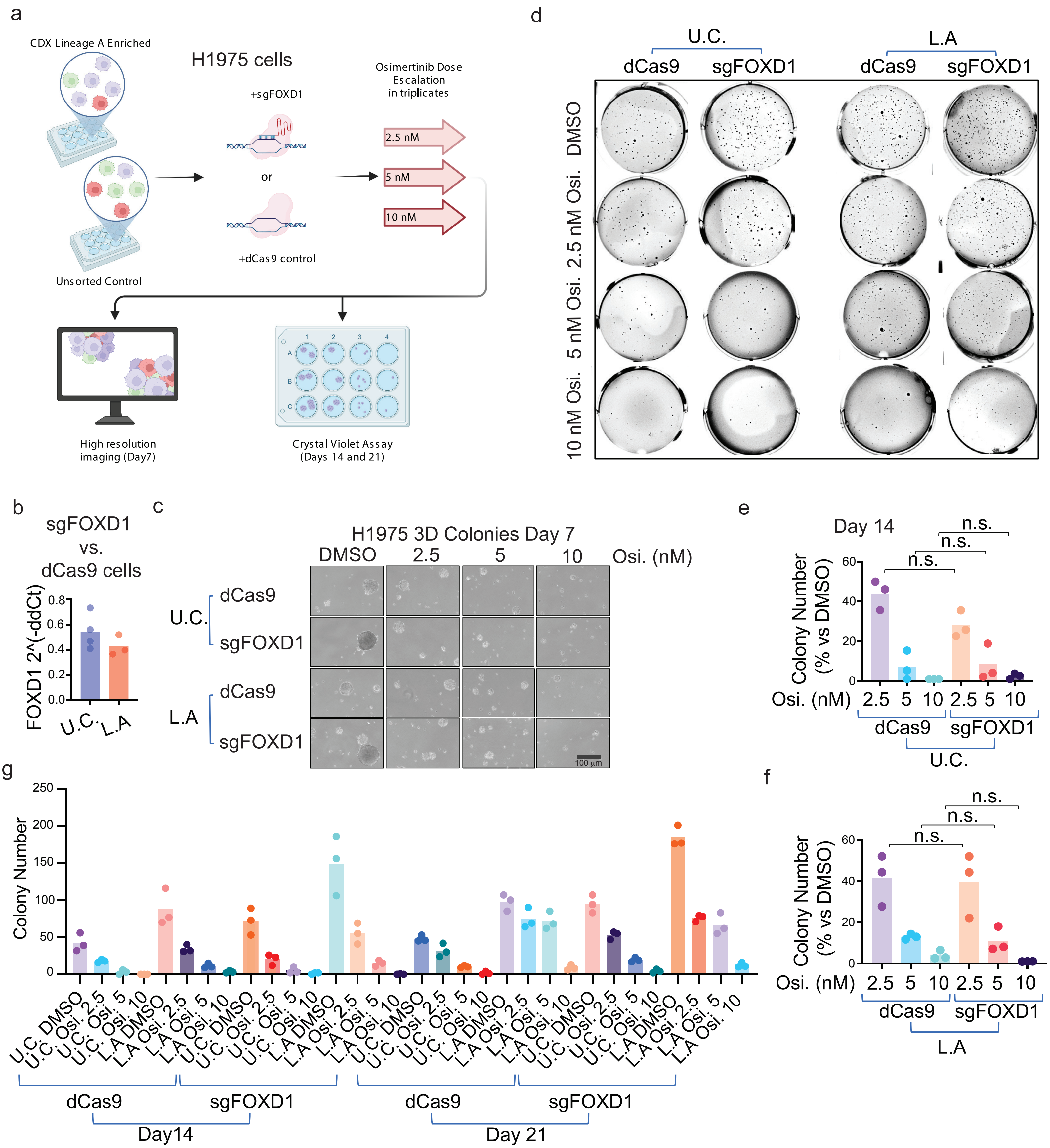

**Supplementary Figure 7**

(a) Workflow of functional tests using CRISPR-Cas9 lentiviral infection with sgRNA targeting FOXD1 protein and 3D-soft agar colony formation assay using lineage A enriched (L.A) or unsorted control (U.C.) H1975 cells treated with a dose escalation of osimertinib (Osi., 2.5-10 nM). Colonies were imaged on day 7, stained with crystal violet on days 14 and 21, and quantified. (b) qPCR analysis attesting successful knockdown of FOXD1 gene using CRISPR-Cas9 lentiviral infection with sgRNA targeting FOXD1 protein in lineage A enriched (L.A) or unsorted control (U.C.) H1975 cells (data normalized to GAPDH and expressed as a ratio to dCas9 control cells). (c) H1975 L.A and U.C. 3-D colony images at day 7 upon dose escalation of osimertinib (2.5-10 nM). (d) Representative images of crystal violet stained-lineage A enriched (L.A) or unsorted control (U.C.) H1975 cells carrying dCas9 or sgFOXD1 were treated with a dose escalation of osimertinib (Osi., 2.5-10 nM). (e, f) H1975 U.C. (e) or L.A (f) dCas9 and sgFOXD1 colonies treated with osimertinib or DMSO control were stained with crystal violet on day 14 and quantified (p-value calculated using Two-way ANOVA with Sidak's multiple comparison test). (g) Total dCas9 or sgFOXD1 colony's number in 3-D experiments with a dose escalation of osimertinib (2.5-10 nM) or DMSO control (crystal violet quantification at day 14 and day 21).

Supplementary Figure 8

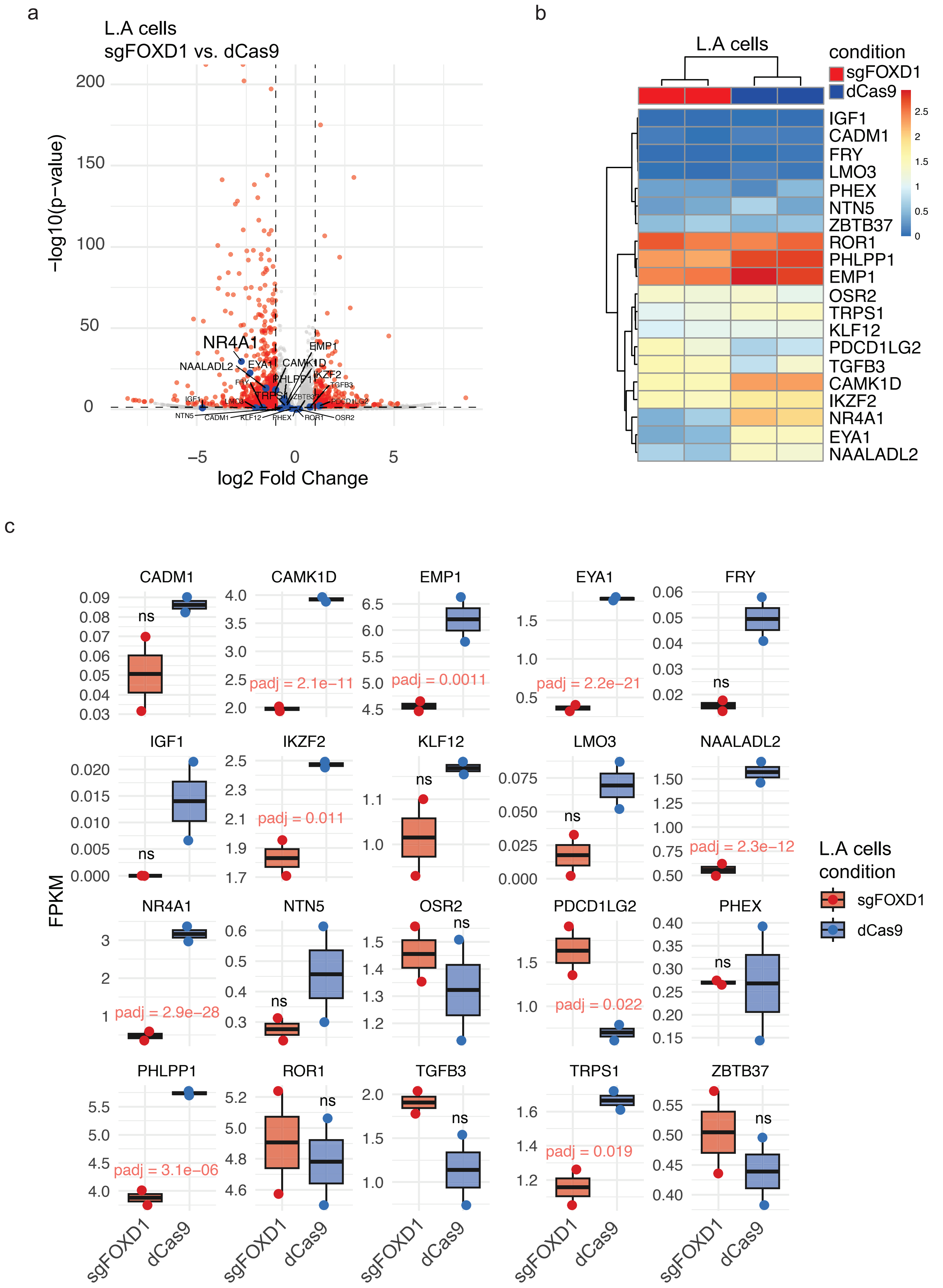

**Supplementary Figure 8**

(a) Volcano plots showing depletion of FREAC4/FOXD1 regulon gene set in RNA sequencing dataset analysis for L.A sgFOXD1 H1975 cells vs. L.A H1975 dCas9 controls (n = 2 samples per group). Blue dots represent FREAC4/FOXD1 regulon genes. Vertical dashed lines represent +/- log2(1.5)FC. (b-c) Heatmap (b) and boxplots (c) visualization of the depletion of FREAC4/FOXD1 regulon gene set in RNA sequencing dataset for L.A sgFOXD1 H1975 cells vs. L.A H1975 dCas9 controls. Adjusted p-values for significance are indicated in the boxplots.

Supplementary Figure 9

a

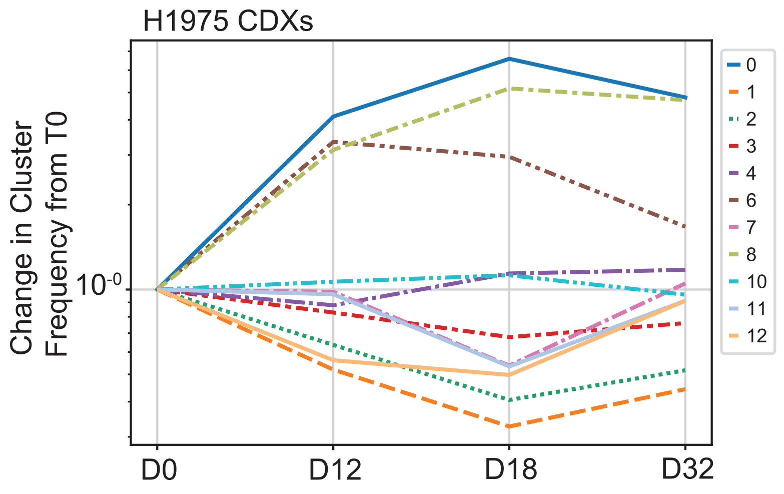


b

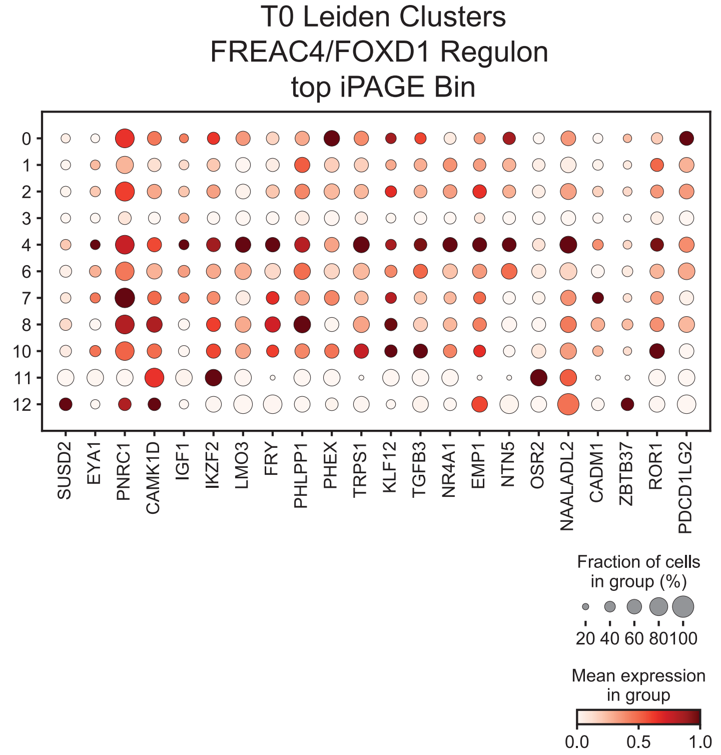


c

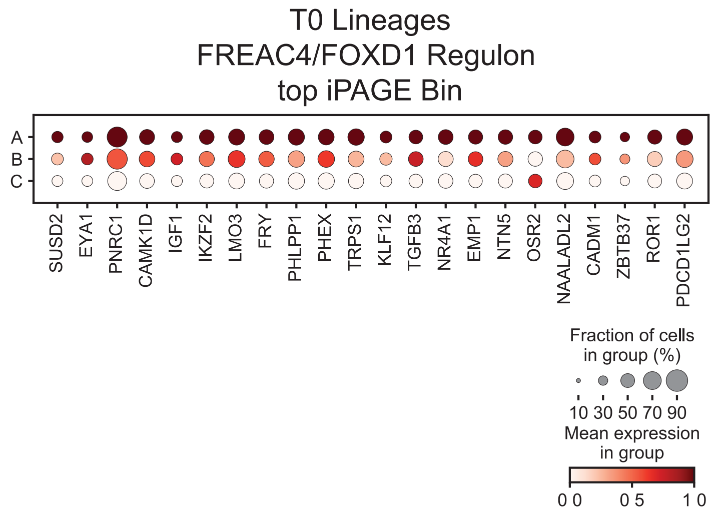

**Supplementary Figure 9**

(a) Leiden clustering of the H1975 CDX data identified thirteen clusters (clusters 0-12); clusters 5 and 9 were identified as dying cells and omitted from further analysis. (b, c) Dot plots with single-cell RNAseq data from H1975 CDXs obtained by checking the expression of the FREAC4 regulon genes from the top iPAGE bin in each of the eleven Leiden clusters (b) and in lineage A-C populations identified by lineage tracing (c).
